# Heterogeneity and targeted therapy-induced adaptations in lung cancer revealed by longitudinal single-cell RNA sequencing

**DOI:** 10.1101/2019.12.08.868828

**Authors:** Ashley Maynard, Caroline E. McCoach, Julia K. Rotow, Lincoln Harris, Franziska Haderk, Lucas Kerr, Elizabeth A. Yu, Erin L. Schenk, Weilun Tan, Alexander Zee, Michelle Tan, Philippe Gui, Tasha Lea, Wei Wu, Anatoly Urisman, Kirk Jones, Rene Sit, Pallav K. Kolli, Eric Seeley, Yaron Gesthalter, Daniel D. Le, Kevin A. Yamauchi, David Naeger, Nicholas J. Thomas, Anshal Gupta, Mayra Gonzalez, Hien Do, Lisa Tan, Rafael Gomez-Sjoberg, Matthew Gubens, Thierry Jahan, Johannes R. Kratz, David Jablons, Norma Neff, Robert C. Doebele, Jonathan Weissman, Collin M. Blakely, Spyros Darmanis, Trever G. Bivona

**Affiliations:** Chan-Zuckerberg Biohub San Francisco, CA; Department of Medicine, University of California, San Francisco, CA; University of California, San Francisco, Helen Diller Family Comprehensive Cancer Center, San Francisco, CA; Lowe Center for Thoracic Oncology, Dana-Farber Cancer Institute; Boston, MA; Department of Cellular and Molecular Pharmacology, University of California, San Francisco, CA; Department of Medicine, University of Colorado Anschutz Medical Campus, Aurora, CO; Department of Biomolecular Engineering, University of California, Santa Cruz, CA; Department of Pathology University of California, San Francisco, CA; Department of Radiology and Biomedical Imaging; University of California, San Francisco, CA; Denver Health Medical Center, Denver, CO; Department of Radiology, University of Colorado, Aurora, CO; Department of Surgery, University of California, San Francisco, CA; Howard Hughes Medical Institute, University of California, San Francisco, CA

**Author notes:** These authors contributed equally (Ashley Maynard, Caroline E. McCoach). These authors contributed equally (Julia K. Rotow, Lincoln Harris).

**Keywords:** single cell RNA sequencing, NSCLC, EGFR, ALK, immune microenvironment, mutation, heterogeneity, landscape

## Abstract

Lung cancer, the leading cause of cancer mortality, exhibits heterogeneity that enables adaptability, limits therapeutic success, and remains incompletely understood. Single-cell RNA sequencing (scRNAseq) of metastatic lung cancer was performed using 44 tumor biopsies obtained longitudinally from 27 patients before and during targeted therapy. Over 20,000 cancer and tumor microenvironment (TME) single-cell profiles exposed a rich and dynamic tumor ecosystem. scRNAseq of cancer cells illuminated targetable oncogenes beyond those detected clinically. Cancer cells surviving therapy as residual disease (RD) expressed an alveolar-regenerative cell signature suggesting a therapy-induced primitive cell state transition, whereas those present at on-therapy progressive disease (PD) upregulated kynurenine, plasminogen, and gap junction pathways. Active T-lymphocytes and decreased macrophages were present at RD and immunosuppressive cell states characterized PD. Biological features revealed by scRNAseq were biomarkers of clinical outcomes in independent cohorts. This study highlights how therapy-induced adaptation of the multi-cellular ecosystem of metastatic cancer shapes clinical outcomes.

## Introduction

Heterogeneity is a property of many biological systems and diseases such as cancer. Biological plasticity in cancer cells is one form of heterogeneity that allows for early adaptation to treatment and limits the success of precision approaches for cancer treatment (Yuan et al., 2019; Xue et al., 2017). In addition to cancer-cell intrinsic heterogeneity, cells within the tumor microenvironment (TME) further contribute to tumor heterogeneity in a cancer-cell extrinsic manner. While these tumor compartments and tumor heterogeneity have been characterized in many cancer subtypes (Gerlinger et al., 2012; Alexandrov et al., 2013; Lawrence et al., 2013; Brannon et al., 2014; Lee et al., 2014; Vignot et al., 2015; Hata et al., 2016), our understanding of how these properties evolve and interact longitudinally in response to systemic treatment remains incomplete, particularly in metastatic tumors.

Many oncogene-driven cancers such as those with oncogenic alterations in *EGFR, ALK, ROS1*, and *BRAF* are treated with targeted therapies against the cognate oncoprotein or signaling pathway. This has led to improvements in clinical outcomes of metastatic solid cancers such as lung cancer and melanoma as well as hematologic malignancies (Flaherty et al., 2012; Mok et al., 2009; Shaw et al., 2013). A paradigm for molecular therapeutics and the study of tumor evolution is lung cancer, the leading cause of cancer mortality worldwide (Siegel et al., 2018), with non-small cell lung cancer (NSCLC) constituting the major subtype (Swanton and Govindan, 2016). Despite the success of targeted therapy in cancers such as NSCLC, tumors typically respond incompletely in the form of residual disease that is a prelude to tumor progression due to therapy-induced tumor evolution. Bulk tumor sampling after tumor progression on targeted therapy has identified resistance mechanisms and demonstrated that tumors become increasingly molecularly heterogeneous after therapy progression (Blakely et al., 2017a; Camidge et al., 2014; McCoach et al., 2018; Rotow and Bivona, 2017).

Single-cell RNA sequencing (scRNAseq) is one established approach to dissect the heterogeneity of complex biological systems (Chung et al., 2017; Darmanis et al., 2017; Tirosh et al., 2016). Comparison of single-cell transcriptional profiles of individual cells at different treatment timepoints could lead to deeper insight into the evolution of cell states of both cancer cells and TME cells occurring during treatment. There is currently a paucity of single-cell transcriptome studies that sample metastatic solid malignancies and prior scRNAseq studies of metastatic disease largely focused on individual treatment timepoints (Chung et al., 2017a; Darmanis et al., 2017; Lambrechts et al., 2018a; Patel et al., 2014a; Tirosh et al., 2016a; Wang et al., 2019; Zhang et al., 2019a). This is due, in part, to challenges associated with obtaining high-quality biopsies from metastatic tumors, instead of the larger tumor specimens that can be obtained from surgically resected earlier-stage disease. Obtaining multiple tumor biopsies longitudinally at different timepoints during systemic therapy of patients, individually and as a cohort, with metastatic solid tumors is an additional challenge to overcome to allow for the study of tumor adaptation to systemic therapy.

Beyond the analysis of tumor biopsies obtained both before treatment and at tumor progression, analysis of advanced-stage tumor samples that reflect the residual disease (*RD*) state during therapy could reveal clinically relevant biological events that permit cancer cell persistence. Developing a deeper understanding of evolving cell-state changes during treatment, and particularly at the RD state, is essential to improve the durability and magnitude of response to precision therapies for cancer. RD sample analysis could help identify therapeutic targets in cancer and TME cells for exploitation in a window-of-opportunity during therapy to preempt the subsequent evolution to absolute drug resistance.

We performed scRNAseq analyses on human NSCLC tumor biopsies collected at different treatment states. Our goal was to help elucidate cancer cell phenotypic plasticity and the contribution and dynamic nature of the TME during therapy that individually, and collectively, promote RD and subsequent progressive disease during targeted therapy. To accomplish this, we instituted a unique clinical tumor rebiopsy protocol wherein advanced-stage NSCLC samples were obtained from patients longitudinally before systemic targeted therapy (*treatment naïve, TN*), at the residual disease state during treatment response (*residual disease, RD*) and upon the subsequent establishment of acquired drug resistance (*progression, PD*). We developed a custom tissue processing and analytic framework adapted to the relatively small and challenging advanced-stage lung tumor biopsy samples to: 1) evaluate the expressed mutational landscape of tumor samples, 2) characterize transcriptional gene signatures unique to different treatment timepoints, 3) investigate the properties of the TME as it evolves during treatment, and 4) describe the interplay between cancer cells and the TME during targeted therapy.

## Results

### Longitudinal scRNAseq analysis of advanced-stage NSCLC during targeted therapy

We used scRNAseq to profile 44 samples (41 lung adenocarcinomas, 1 squamous cell carcinoma and 2 normal adjacent tissues) (Figure 1A), corresponding to 27 individual patients. We used a rapid processing workflow customized to isolate viable single cells primarily from fine needle aspirates (FNA), core needle biopsies, and thoracentesis samples in addition to standard surgical resections (Figure 1B). Tumor biopsy samples were collected and immediately processed before and during systemic targeted therapy (Figure 1C, Supplemental Table 1).

**Table 1.** Table of common and unique features in different treatment timepoints and possible therapeutic approaches.

| ID | Feature | Therapeutic approach | References |
| --- | --- | --- | --- |
| 1 | Multiple targetable mutations | Combination targeted therapy | (McCoach and Bivona, 2019) |
| 2 | Invasion pathways | Targeted inhibition | (Rahman et al., 2017b; Zhang et al., 2016, 2019b) |
| 3 | Increased immunostimulatory T-cells | Immune system modulation | (Ha et al., 2019; Jenkins et al., 2018; Souza-Fonseca-Guimaraes et al., 2019; Valkenburg et al., 2018) |
| 4 | Alveolar signature | Targeted inhibition | (Nabhan et al., 2018c; Zhang et al., 2019b) |
| 5 | Plasminogen activation signature | Targeted inhibition | (Mahmood et al., 2018; Zhang et al., 2019b) |
| 5 | SERPINE1 signature | Targeted inhibition and immune modulation | (Placencio and DeClerck, 2015) |
| 6 | Gap Junction signature | Targeted inhibition | (Wu and Wang, 2019; Mulkearns-Hubert et al., 2019) |
| 7 | Loss of Tumor suppressors | Targeting acquired vulnerabilities | (Ding et al., 2019) |
| 8 | Increased pro-inflammatory chemokines | Immune system modulation | (Tokunaga et al., 2018) |
| 9 | Increased Tregs | Immune system modulation | (Tanaka and Sakaguchi, 2019) |
| 10 | Kyurenine signature | Targeted inhibition and immune modulation | (Labadie et al., 2019) |

**Fig. 1.**
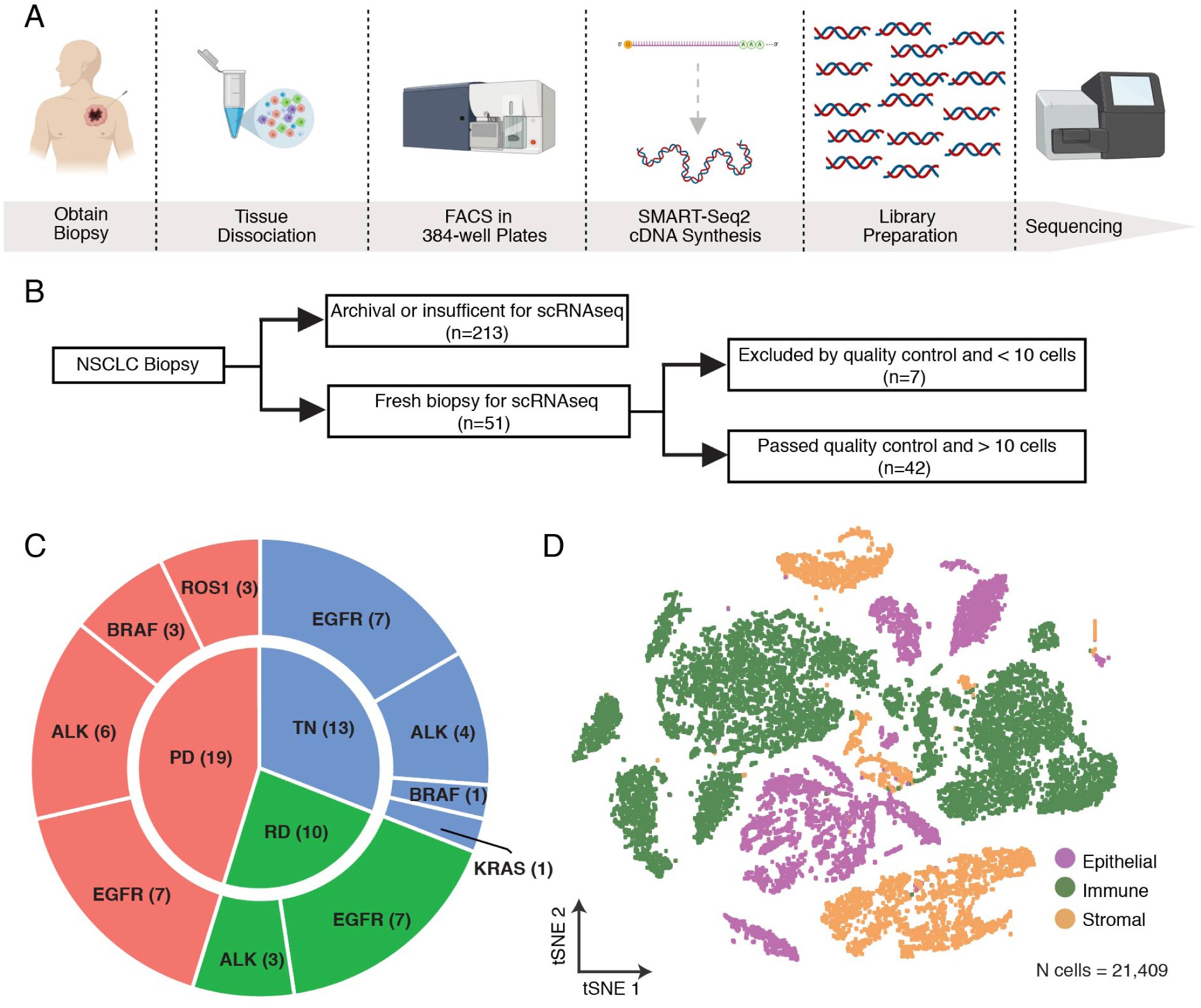
Patient characteristics and experimental overview. Tissue processing pipeline for scRNAseq. Patient samples were first mechanically, and enzymatically disaggregated and then single cells were sorted into microtiter plates using FACS. cDNA synthesis was performed using the Smart-seq2 protocol and libraries were sequenced on Illumina platforms. (B) Consort diagram of sample collection. A total of 51 biopsies were processed. Only samples with cells that passed quality control were used (n = 42). (C) Circle plot illustrating the clinically identified oncogenic driver (outer circle) and treatment timepoint (inner circle) for each sample. (D) t-SNE plot of all 21,409 cells colored by their cellular identity as inferred from gene expression profiling (Epithelial cells (n= 5,109, Immune Cells (n=12,077), Stromal Cells (n=4,223).

Gene expression profiles of 21,409 cells were retained after quality control filtering (Supplemental Figure 1A). Following gene expression normalization, we performed principal component analysis (PCA) using a set of variably expressed genes across all cells. We clustered cells using graph-based clustering on the informative principal component space (npcs=20). The resulting cell clusters were annotated as immune cells, stromal cells (fibroblasts, endothelial cells, and melanocytes), and epithelial cells (Figure 1D) through gene expression analysis of established marker genes (Lambrechts et al., 2018; Schiller et al., 2019; Tabula Muris Consortium et al., 2018; Treutlein et al., 2014) (Supplemental Table 2). Epithelial cells (n=5,109), were subset and re-clustered into 31 discrete epithelial clusters (Supplemental Figure 1B).

### Clustering-based copy number variation resolves cancer from non-cancer epithelial cells

Given the known association between cancer and large-scale chromosomal alterations, we utilized an established analytical framework to infer copy number variation (CNV) from RNA expression to classify epithelial cells as either cancer or non-cancer (Patel et al., 2014; Puram et al., 2017; Tirosh et al., 2016; Venteicher et al., 2017; https://github.com/broadinstitute/inferCNV). Compared to fibroblasts and endothelial cells, which were used as controls, cancer cells displayed larger excursions from relative expression intensities in multiple regions of the genome (Supplemental Figure 1C). Following identification of cancer and non-cancer epithelial cells, non-cancer epithelial cell clusters (n=13) were further annotated as either pulmonary alveolar type 2 (AT2), pulmonary alveolar type 1 (AT1), hepatocytes, club cells, basal cells, ionocytes or stromal cells (Supplemental Figure 1D).

As noted by others (Zhang et al., 1997), we found that these cancer cells expressed an elevated number of unique genes compared to non-cancer cells (Supplemental Figure 1E). This difference in the number of uniquely expressed genes was not explained by sequencing depth (Pearson correlation = 0.17).

All cancer cells (n=3,620) were subset and re-clustered, resulting in 24 unique clusters across 38 total samples (Supplemental Figure 2A, 2B). For each cluster, we calculated patient occupancy, defined as the number of cells of the highest contributing individual patient over the total number of cells for that cluster for both non-cancer and cancer epithelial cells (Supplemental Figure 2C). The majority of the cancer cell clusters are patient specific, having high patient occupancy scores, similar to findings reported in other studies (Chung et al., 2017; Darmanis et al., 2017; Puram et al., 2017; Tirosh et al., 2016). Conversely, non-cancer cell types exhibited lower patient occupancy, with multiple patients contributing cells to each cluster (Supplemental Figure 2C). Thus, patient-specific malignant cell clustering reflects the unique molecular signatures of an individual patient’s tumor rather than potential technical issues.

**Fig. 2.**
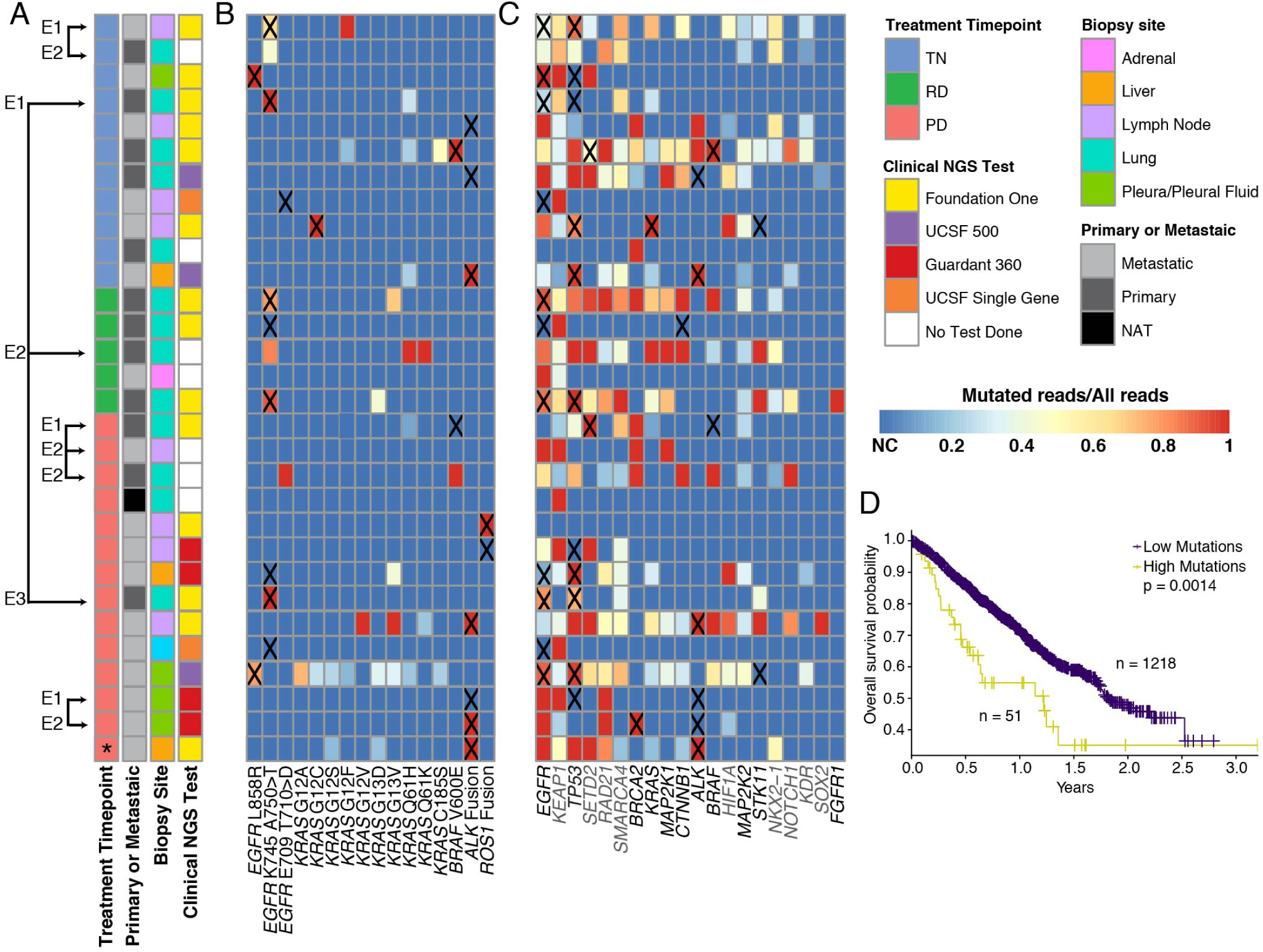
scRNAseq infers patient mutational status and reveals a complex mutational landscape in cancer cells. (A) Clinical characteristics of the 30 NSCLC samples from which oncogenic drivers and/or COSMIC tier 1 genes were identified. Columns indicate treatment timepoint (TN, RD, PD), primary or metastatic sample origin, biopsy site, and clinical NGS test performed. Longitudinal samples from the same patient are indicated by chronological encounter numbers (e.g. E1, E2) and grouped with arrows. (B) Cancer cell mutational landscape for each patient sample as determined by scRNAseq represented as a heatmap. Color indicates the number of mutant reads for each genomic region and sample divided by the total number of reads for that region in that sample. The expected oncogenic driver, as determined by a clinical NGS assay, is marked with an “X”. NC-No Coverage for the specific sample. (C) Mutational landscape of COSMIC tier 1 genes. Color indicates the number of mutant reads for each genomic region and sample divided by the total number of reads for that region in that sample. Genes names in black indicate inclusion in all clinical NGS assays (aside from the UCSF single gene test), whereas gene names in grey are included in only select assays (Supplemental Table 2). D) Kaplan-Meier plot showing overall survival of 1269 NSCLC patients within the MSK-Impact dataset. Patients were stratified by high (>=2) and low (<2) mutations from the 141 mutations that are found in both the MSK-Impact dataset and panel (C).

### scRNA-seq analysis reveals a rich complexity of expressed gene alterations in cancer cells

Additional genetic alterations detected by bulk tumor DNA-based analysis can co-exist with a primary targetable oncogenic “driver” alteration (e.g. oncogenic *EGFR, ALK, BRAF, ROS1, KRAS* and *MET*) and may help promote tumor progression and limit therapy response (Blakely et al., 2017; Kim et al., 2019; Scheffler et al., 2019; Yang et al., 2019). We queried scRNAseq transcripts from each cancer cell to identify somatic alterations, namely single nucleotide polymorphisms (SNPs), insertions/deletions (indels), and gene fusions (Figure 2A, 2B, 2C). We identified 17 of 38 tumor biopsy samples that contained cancer cells harboring the clinically-identified oncogenic driver (Figure 2B, Supplemental Table 3), consistent with the known potential drop-out occurrence in scRNAseq analyses (Kharchenko et al., 2014). Within these 17 samples in which we found at least one cell harboring the clinically actionable mutation, there were 7 samples in which at least one other cell harbored an additional oncogenic alteration not detected by clinical-grade testing of a tumor sample from the same patient (i.e. occult genetic alterations) (Figure 2B). One such example from our data is illustrated by sample LTS47. This tumor was determined to harbor an *EML4-ALK* oncogenic gene rearrangement by clinical-grade bulk DNA analysis. However, scRNAseq additionally revealed that this sample contained cancer cells harboring *KRAS* G13D and *KRAS* G12C occult mutations; neither population of *KRAS* mutant cells showed evidence of the *ALK* gene rearrangement (Supplemental Figure 2D). This sample was obtained from the patient after multiple lines of therapy which likely allowed for evolution of multiple mechanisms of resistance, including the 2 different oncogenic forms of *KRAS*, which are known mechanisms of resistance to *ALK* inhibitor treatment (Doebele et al., 2012; Hrustanovic and Bivona, 2015; Shaw and Engelman, 2013).

We also queried scRNAseq transcripts for mutations from the COSMIC (Catalogue of Somatic Mutations In Cancer) lung adenocarcinoma tier 1 mutations (Supplemental Table 2), (Forbes et al., 2017; Shihab et al., 2015), focusing on mutations that are classified as ‘pathogenic’. Many of the mutations we identified had not been previously reported by the clinical-grade assay conducted on the patient’s tumor despite having been included in the clinical panel (Figure 2C, Supplemental Tables 2 and 3). Though this may reflect the difference in biopsy technique or tumor clonality at the time of clinical testing, these results also demonstrate that clinical-grade bulk DNA-based testing may underestimate tumor heterogeneity. To assess the clinical outcomes of patients harboring multiple oncogenic alterations and to determine the broader translational impact of our findings, we utilized the MSK-Impact NSCLC dataset, which contains the clinical and sequencing data for over 800 advanced-stage NSCLC patients (Zehir et al., 2017). Those patients with greater than or equal to 2 mutations from the tier one COSMIC mutations found in the scRNAseq dataset (mutation high), had significantly lower overall survival (p < 0.01) compared to those with less than 2 mutations (mutation low) Patients with more than two cosmic tier 1 mutations (Figure 2D). Thus, scRNAseq analysis can complement current clinical-grade bulk DNA-based assays to provide increased granularity into cancer cell genomic heterogeneity and provides insight into the mutational landscape that is expressed at the RNA level in cancer cells of advanced-stage tumors, with clinical implications.

### Transcriptional differences between TN and RD cancer cells detected by scRNAseq analysis reveal cell state-specific biological programs

We hypothesized that defining the biological programs activated in cancer cells during therapy response may identify signaling pathways that promote adaptation and survival of cancer cells that comprise residual disease during initial treatment. We compared the transcriptional profiles of individual cancer cells obtained from tumor samples from TN to RD (Supplemental Table 4) and focused our attention on the 691 significantly (p<0.05) upregulated genes in RD cancer cells as a proxy for evidence of pathway activation. We found individual genes associated with specific cancer-associated pathways such as survival, cell growth, invasion and metastasis (Supplemental Table 5). As a control and consistent with the expectation that during targeted treatment cancer cells surviving and persisting drug therapy are generally less proliferative (Sharma et al., 2010; Zhu et al., 2001), we found that RD cancer cells expressed decreased proliferation marker genes compared to both TN and PD (Supplemental Figure 3A) (Hsiao et al., 2019).

**Fig. 3.**
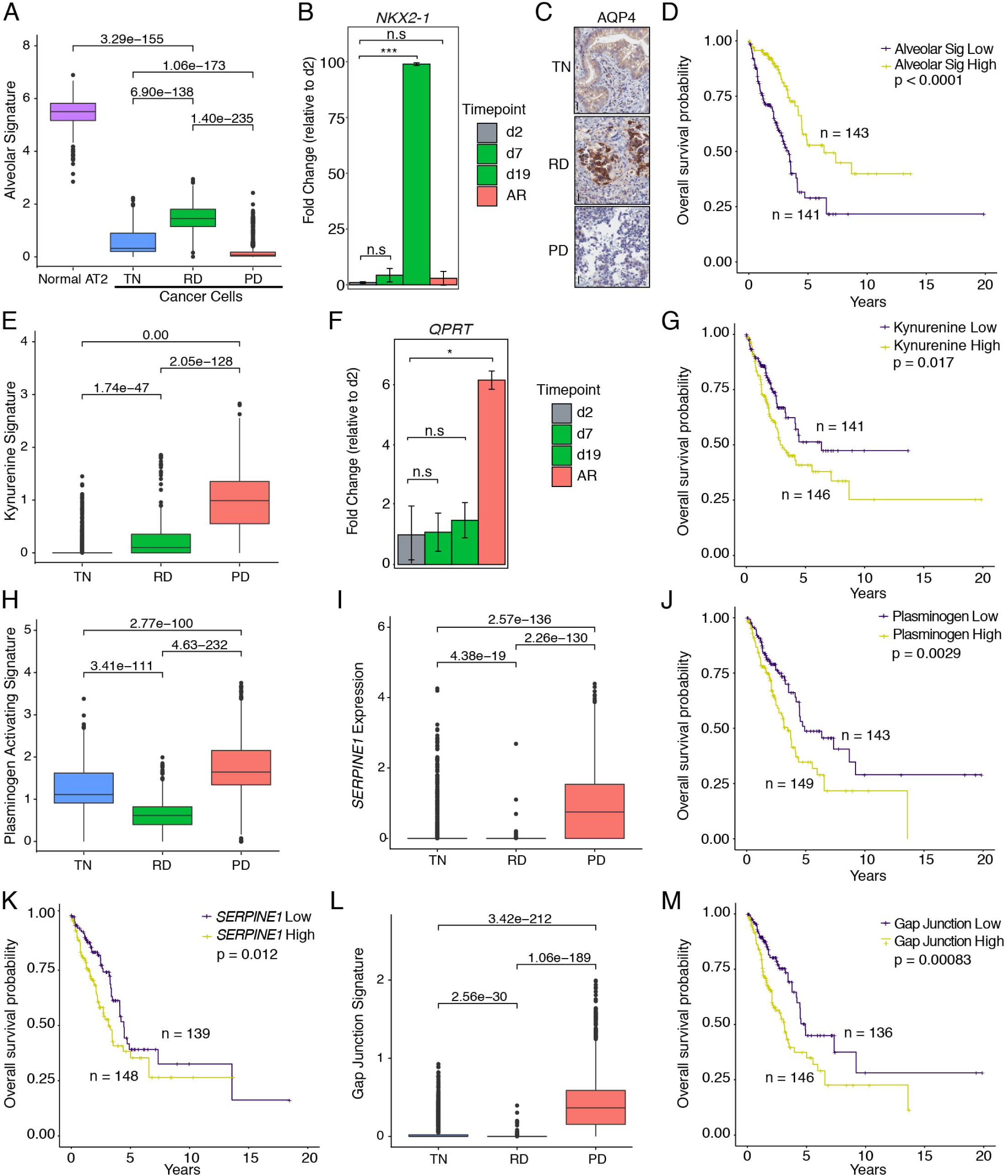
Differential gene-expression analysis between treatment timepoints reveals treatment stage specific transcriptional signatures. (A) Boxplots showing the expression level of the alveolar signature across different treatment timepoints as well as non-cancerous AT2 cells from our cohort. (B) Fold change expression of *NKX2-1* as quantified by RT-PCR in PC9 cells after treatment with vehicle (2 days), osimertinib at 7 days (RD), 19 days (RD) and at the acquired resistance state for this cell line, day 70 (see methods) (AR), *** indicates p < 0.001. (C) Representative IHC images of TN, RD and PD tumor tissue sections stained for AQP4 demonstrating increased expression at the RD timepoint. Scale bars correspond to 50um. See Supplemental Figure 3D, E for validation cohort. (D) Kaplan-Meier plot of the relationship between the alveolar signature and patient OS within the TGCA dataset. Patients were stratified by high (n=143) and low (n=141) signature expression. (E) Box plots showing the expression levels of the kynurenine signature expression across different treatment timepoints. (F) Fold change expression of QPRT as quantified by RT-PCR in PC9 cells after treatment with osimertinib as in (B) (see methods) (AR), * indicates p < 0.05. (G) Kaplan-Meier plot of the relationship between the kynurenine signature and patient OS within the TGCA dataset. Patients were stratified by high (n=146) and low (n=141) signature expression. (H, I) Box plots showing the expression levels of the plasminogen activation, *SERPINE1* pathway signatures across different treatment timepoints. (J, K) Kaplan-Meier plots of relationships between the plasminogen activating pathway signature and *SERPINE1* and patient OS within the TGCA dataset, respectively. Patients were stratified by high and low signature expression. (L) Box plot showing the expression levels of the gap junction signature across treatment timepoints. (M) Kaplan-Meier plots of relationships between the gap junction signature and patient OS within the TGCA dataset. Patients were stratified by high and low signature expression.

Interestingly, we identified an alveolar cell gene expression signature composed of 17 established gene markers of alveolar cells (Vieira Braga et al., 2019; Wade et al., 2006) that showed significantly (p< 0.0001) increased expression in RD versus TN timepoints (Figure 3A, Supplemental Figure 3B, Supplemental Table 2). Alveolar cells are partitioned into alveolar type 1 (AT1) and type 2 (AT2) subtypes and form the lining of the lung alveoli. AT2 cells produce surfactants and can act as stem-like progenitor cells (Nabhan et al., 2018) which become active and proliferate in the setting of diverse types of lung injury. AT2 cell are also suspected to be the cell of origin in oncogene-driven lung cancers (Desai et al., 2014; Hanna and Onaitis, 2013; Nabhan et al., 2018). AT1 cells are the dominant population in alveoli and mediate gas exchange and, when injured or dying, can release proliferation and regenerative signals (Desai et al., 2014). The alveolar signature we detected in the cancer cells at RD includes increased expression of both AT1- and AT2-associated genes (Supplemental Table 2), including *AQP4, SFTPB/C/D, CLDN18, FOXA2, NKX2-1* and *PGC* for AT2 cells (Desai et al., 2014; Liu et al., 2003; Nabhan et al., 2018; Wade et al., 2006; Xu et al., 2016; Zhou et al., 2018) and *AGER* and *HOPX* for AT1 cells (Nabhan et al., 2018a; Serveaux-Dancer et al., 2019) (Supplemental Figure 3B). Additional analysis demonstrated that the alveolar cell state we identified in cancer cells was not derived from misannotated non-cancer alveolar cells within our cancer cell populations (Supplemental Figure 3C).

We validated the activation of the alveolar cell signature at RD using orthogonal approaches. First, we used an established preclinical model consisting of patient-derived EGFR-mutant NSCLC cells (PC9) (Lee et al., 1985) to develop analogs of the TN, RD, and PD clinical states and measured the expression of a subset of the alveolar cell state genes (by RT-PCR) detected at RD in the clinical tumor samples. Cells were treated with vehicle (DMSO) or a standard EGFR inhibitor (osimertinib) and sampled under control conditions (day 2 in DMSO), during RD (days 7 and 19), and at PD (day 70, acquired resistance). We found significantly increased expression of *NKX2-1*, a hallmark alveolar cell signature gene upregulated in the RD clinical samples, at days 7 and 19 compared to control and day 70 (Figure 3B). This demonstrates that our findings from the scRNAseq analysis can be reproduced under controlled conditions *in vitro*. Furthermore, immunohistochemistry (IHC) analysis showed induction of AQP4 protein expression, another marker of the alveolar cell signature, at the plasma membrane of RD clinical samples compared to both TN and PD clinical samples (Figure 3C, Supplemental Figure 3D-E). The collective data support key scRNAseq findings arising from the clinical tumor biopsies.

We next determined whether the alveolar cell signature was more broadly clinically-relevant by examining whether it is a biomarker of patient survival in the TGCA lung adenocarcinoma bulk RNAseq dataset generated by the TCGA Research Network (https://www.cancer.gov/tcga, Cancer Genome Atlas Research Network et al., 2013). We found a significant (p<0.0001) association between high expression of our alveolar cell type signature and improved overall survival (OS) (Figure 3D). These collective findings support the assertion that there is a distinct alveolar cell type gene expression signature characterizing RD cancer cells that is associated with improved patient survival. A plausible model is that our identified alveolar signature reflects a cell injury and repair signal in which the machinery of injury-response in malignant AT1/2 cells activates a regenerative signaling pathway and transition to a more primitive cell state. This could serve to aid repair from injury from cell death during treatment to support cancer cell persistence, while at the same time generating a less aggressive malignant state. This is consistent with notion that RD represents a “persister” cell state observed in preclinical models of slow-cycling cancer cells that survive without rapid proliferation (as in Supplemental Figure 3A), as a prelude to the onset of aggressive tumor progression upon absolute drug resistance (Hata et al., 2016).

The molecular details of the alveolar and cell injury repair signature are notable. In our RD cohort, WNT/β-catenin-associated pathway genes *AFF3, SUSD2*, and *CAV1* exhibited increased expression (Supplemental Table 5). *AFF3* and *SUSD2* are activated downstream targets of the WNT pathway (Lefèvre et al., 2015; Umeda et al., 2018; Xu et al., 2018) while *CAV1* can promote nuclear localization of β-catenin (*CTNNB1*) and transcriptional activation of the WNT/β-catenin pathway (Yu et al., 2014). In NSCLC, the WNT/β-catenin signaling pathway contributes to tumorige-nesis (Juan et al., 2014; Nakayama et al., 2014; Pacheco-Pinedo et al., 2011), repair, and regeneration after cell injury (Huch et al., 2013; Tammela et al., 2017). The self-renewal and injury response in AT2 cells specifically can utilize the WNT/β-catenin signaling pathway (Nabhan et al., 2018b; Stewart, 2014). Additionally, in *EGFR*-mutant NSCLC the WNT/β-catenin pathway may limit EGFR inhibitor response and can lead to survival of a persister cell population during EGFR inhibitor therapy *in vitro* (Arasada et al., 2018; Blakely et al., 2017; Casás-Selves et al., 2012; Nakayama et al., 2014). Overall, the RD state is characterized by signals of cellular injury and survival which act, in part, through the WNT/β-catenin pathway, which may be therapeutically targetable (Krishnamurthy and Kurzrock, 2018).

### Transcriptional differences between TN and PD cancer cells reveal immune modulation and cellular invasion as key features of cancer progression

When comparing cancer cells from TN and PD samples, we found 958 differentially expressed genes with higher expression in PD cancer cells (Supplemental Table 4). Within those genes, we identified genes involved in the kynurenine pathway and multiple genes and pathways associated with tumor invasion, metastasis, cell viability, and inflammation (Supplemental Table 5). We observed a significant (p < 0.0001) increase in the expression of *IDO1, KYNU*, and *QPRT*, which are each involved in the kynurenine pathway, in the PD versus TN cancer cells (Figure 3E, Supplemental Figure 3F, Supplemental Table 2). The kynurenine pathway metabolizes tryptophan generating catabolite intermediates that are further processed to replenish cellular NAD+ and support immunomodulatory roles (Triplett et al., 2018; Zhai et al., 2018). Expression of *IDO1, KYNU*, and *QPRT* can result in immunosuppressive behavior (Triplett et al., 2018). Our data indicate that cancer cells within PD tumors may inhibit the activity of the immune system during targeted therapy. The identification of this pathway as a mediator of immune suppression within PD tumors has important potential therapeutic implications, as IDO1 is known to be upregulated in many cancers (Cheong and Sun, 2018; Hornyák et al., 2018; Liu et al., 2018). Multiple clinical trials have attempted to block this pathway using IDO1 inhibitors as a monotherapy as well as in combination with immune checkpoint inhibitors or hormone therapy (Ricciuti et al., 2019), albeit with limited success. *QPRT* also exhibited increased expression specifically at day 70 of EGFR inhibitor treatment (i.e. acquired resistance and the analog of clinical PD) of our *in vitro* model using PC9 cells (Figure 3F), reinforcing the assertion that this pathway is indicative of cancer progression under the selective pressure of treatment.

To further demonstrate the clinical relevance of the kynurenine pathway, we again used the TCGA lung adenocarcinoma RNAseq dataset. Increased expression of the kynurenine pathway signature was a biomarker of worse OS (p = 0.017) (Figure 3G). This is consistent with the notion that activation of this pathway leads to immunosuppression and an inability of the immune system to effectively surveil and eradicate cancer cells.

### Longitudinal scRNAseq profiles of cancer cells change from RD to PD

We compared cancer cells from RD and PD patient samples to elucidate the differences that occur during the outgrowth of PD from RD and found a total of 2011 genes which had significantly (p<0.001) increased expression in either RD or PD (NRD=874, NPD=1,137) (Supplemental Table 4). Among the differentially overexpressed genes at RD were genes associated with the alveolar cell signature, cell growth, differentiation, and cell motility (Supplemental Table 5). RD cancer cells overexpress surfactant genes (*SFTPB/C/D* and *SFTA3*) which are part of the alveolar cell signature (Figure 3A, Supplemental Figure 3B) (Desai et al., 2014; Treutlein et al., 2014; Wang et al., 2018). Furthermore, RD cancer cells also overexpress the putative tumor suppressor *DLC1*, which is a member of the RhoGAP family of negative regulators of Rho GTPases and can be downregulated in NSCLC (Healy et al., 2008; Yuan et al., 2004). High expression of *DLC1* was associated with an improved survival in NSCLC (Sun et al., 2019). *NKX2-1* and *NFIX* were overexpressed in RD cancer cell and are associated with decreased cell motility (Ge et al., 2018; Rahman et al., 2017a; Winslow et al., 2011). Low expression of *NKX2-1* leads to loss of differentiation and enhanced tumor seeding ability (Winslow et al., 2011). The collective findings arising from this and the previous RD cancer cell analyses suggest that an injury-repair and regenerative cell state may promote cancer cell indolence, increased tumor control, and improved clinical outcomes.

By contrast, PD cancer cells differentially overexpressed genes associated with invasion, cell-to-cell communication, differentiation and immune modulation (Supplemental Table 5). Several genes in the plasminogen activation pathway were significantly overexpressed (*ANXA2, PLAT, PLAUR, PLAU*) (Figure 3H) along with the plasminogen inhibitor *SERPINE1* (PAI1) (p<0.0001, Figure 3I, Supplemental Figure 3G). *ANXA2* and *PLAUR* are the receptor proteins in the plasminogen activation cascade and involved in inflammation, angiogenesis, invasion and metastasis, via degradation of the extracellular matrix (Kubala et al., 2018; Zhu et al., 2017). Signaling is initiated when *ANXA2* or *PLAU* binds to *PLAT* (uTa) or *PLAU* (uPa), respectively. Plasminogen is then degraded to plasmin through the activity of *PLAT* and/or *PLAU* leading to activation of metalloproteinases and degradation of fibrin. *SERPINE1* shows increased expression in a number of cancer subtypes and plays important roles in cell adhesion, invasion, tumor vascularization, radio-resistance, and immunosuppression (Kubala et al., 2018; Zhu et al., 2017). High expression of the plasminogen activation signature correlated with worse OS (p=0.0029) within the TCGA lung adenocarcinoma RNAseq dataset (Figure 3J). Similarly, in this independent dataset high expression of *SERPINE1* was associated with worse OS (p=0.012) (Figure 3K). Interestingly, EGFR inhibitor therapy can induce expression of *SERPINE1* and EGFR mutation positive patients with greater than two-fold induction of *SERPINE1* (PAI1) plasma levels during EGFR inhibitor treatment demonstrate shorter progression free survival (Arasada et al., 2018). Collectively, our longitudinal scRNAseq-based findings shed light on the clinical relevance and potential role of the plasminogen activation cascade in inferior clinical outcomes and targeted therapy resistance.

Additionally, we found several gap junction proteins differentially overexpressed in PD cancer cells compared to RD cancer cells (p<0.0001, Figure 3L, Supplemental Table 2). Gap junction proteins (e.g. connexins) are integral membrane proteins that allow for cytosolic exchange of ions, metabolites and secondary messengers between cells (Aasen et al., 2016; Sinyuk et al., 2018). While some have been identified as tumor suppressors, we found that high expression of *GJB2/3/4/5* (Supplemental Figure 3H) was linked to worse survival in the TCGA lung adenocarcinoma RNAseq dataset (p=0.00083) (Figure 3M). These collective findings suggest a pro-tumor effect not only in our cohort but also in NSCLC more generally.

In summary, within cancer cells we identified a rich complexity of clinically-relevant, expressed mutations that may impact therapy response. Furthermore, evaluation of transcriptional profiles of individual tumor cells longitudinally across different treatment timepoints identified several clinically-relevant cell state changes (Supplemental Figure 3I). The therapy-induced cancer cell adaptions highlighted in our study offer new potential biomarkers and therapeutic strategies for patients with advanced-stage NSCLC.

### Longitudinal scRNAseq analysis of an individual patient’s tumor during treatment

Obtaining consecutive clinical tumor biopsies from individual advanced-stage lung cancer patients before and during treatment is challenging. Nevertheless, we obtained samples from the same primary tumor site from 3 treatment timepoints from a patient (TH226) whose tumor contained a standard EGFR exon 19 deletion oncogenic mutation and was treated with the EGFR inhibitor osimertinib (Supplemental Figure 4A-C). In all 3 biopsies, we identified by scRNAseq RNA expression of the EGFR exon 19 driver mutation in the cancer cells and several other mutations of interest (Supplemental Figure 4D).

**Fig. 4.**
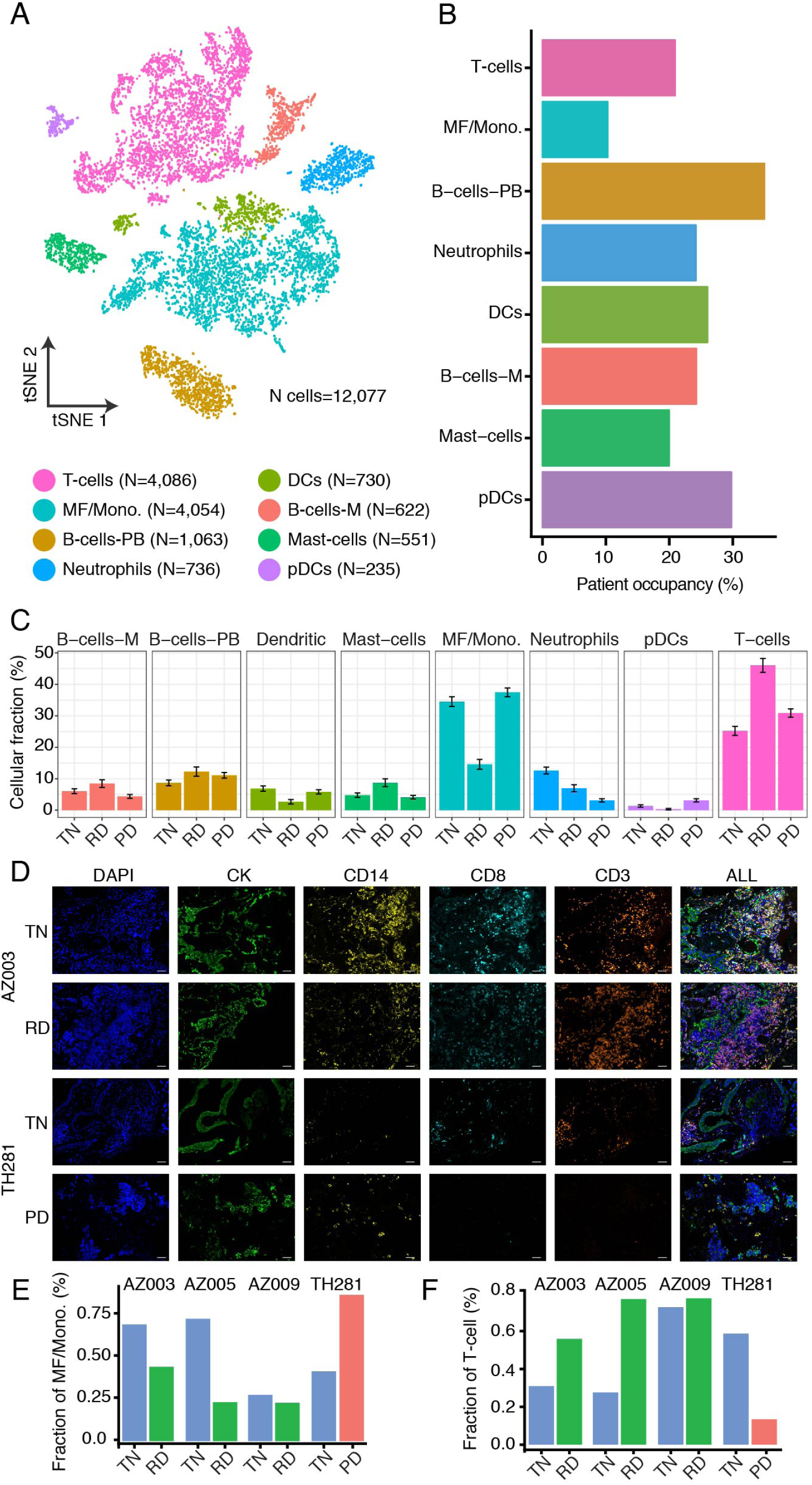
Changes in the composition of the tumor microenvironment within each tumor. (A) t-SNE plot of all immune cells colored by immune cell type. Number of cells for each cell type is provided on the figure legend. (B) Patient occupancy for each immune cell type. (C) Fractional changes for each immune cell type across the three treatment states. Error bars indicate the 95% confidence interval for the calculated relative frequencies. (D) Representative in situ immunofluorescence images of changes from TN to RD and TN to PD in tumor tissue sections from two separate samples at two separate timepoints; AZ003(TN and RD), TH281 (TN and PD). Scale bars correspond to 50 µm. (E and F) Quantification of the fractional changes of macrophages and T-cells, respectively, across treatment timepoints from the images in D and Supplemental Figure 5F.

When comparing TH226 to the rest of scRNAseq dataset, we found overlapping differentially expressed genes and signatures (Supplemental Table 4, Supplemental Figure 4E-H). Intriguingly, we also found numerous genes associated with squamous cell differentiation (*KRT16, KRT14, KRT6A, KRT5, CLCA2, PKP1, ANXA8, DSG3*) overexpressed at PD compared to TN and RD timepoints (p<0.0001, Supplemental Figure 4I, Supplemental Tables 4-5) (Ben-Hamo et al., 2013; Chao et al., 2006; Goodwin et al., 2017). This is particularly interesting given that the patient’s lung tumor biopsy at PD demonstrated a histologic shift to squamous cell carcinoma from that of prior biopsies that showed pure adenocarcinoma histology (Supplemental Figure 4C). Histologic transformation to squamous cell carcinoma is a mechanism of EGFR inhibitor resistance in EGFR-mutant NSCLC (Izumi et al., 2018; Jukna et al., 2016). Thus, scRNAseq has the power to provide a high resolution, gene and pathway level view of biological and histological plasticity that arises during cancer drug treatment.

### Longitudinal inversion of myeloid and lymphoid infiltration within the TME at progressive disease compared to residual disease

We next addressed the evolution of the TME during targeted treatment. The immune cells (n=12,077) were separately clustered and annotated as primary immune cell types (Figure 4A, Supplemental Table 4). In contrast to clusters of cancer cells, which clustered primarily from a single patient sample (Supplemental Figure 2C), immune cell type clusters were each composed of cells derived from multiple different patients and biopsies (Figure 4B). This is consistent with the expectation of finding common immune cell phenotypes across patients and samples. We compared the immune cell composition across all 3 timepoints, expressed as the correlation between fractional immune cell abundance vectors. The immune composition within RD was the most dissimilar from the other two treatment states (r=0.63 versus TN samples, r=0.7 versus PD samples, Pearson’s correlation coefficient) (Supplemental Figure 5A). Across all treatment timepoints, T-cells and macrophages were the dominant cell populations and demonstrated an inversion in relative abundance during tumor response and resistance to treatment, a finding we examined further as described below (Figure 4C). T-cells comprised a larger fraction of all immune cells within the TME at RD compared to TN or PD samples (25% T-cells TN, 46% RD, 32% PD). Macrophage infiltration followed the inverse pattern, with a decrease in macrophages at RD compared to TN and PD (34% Macrophages TN, 15% RD, 37% PD).

**Fig. 5.**
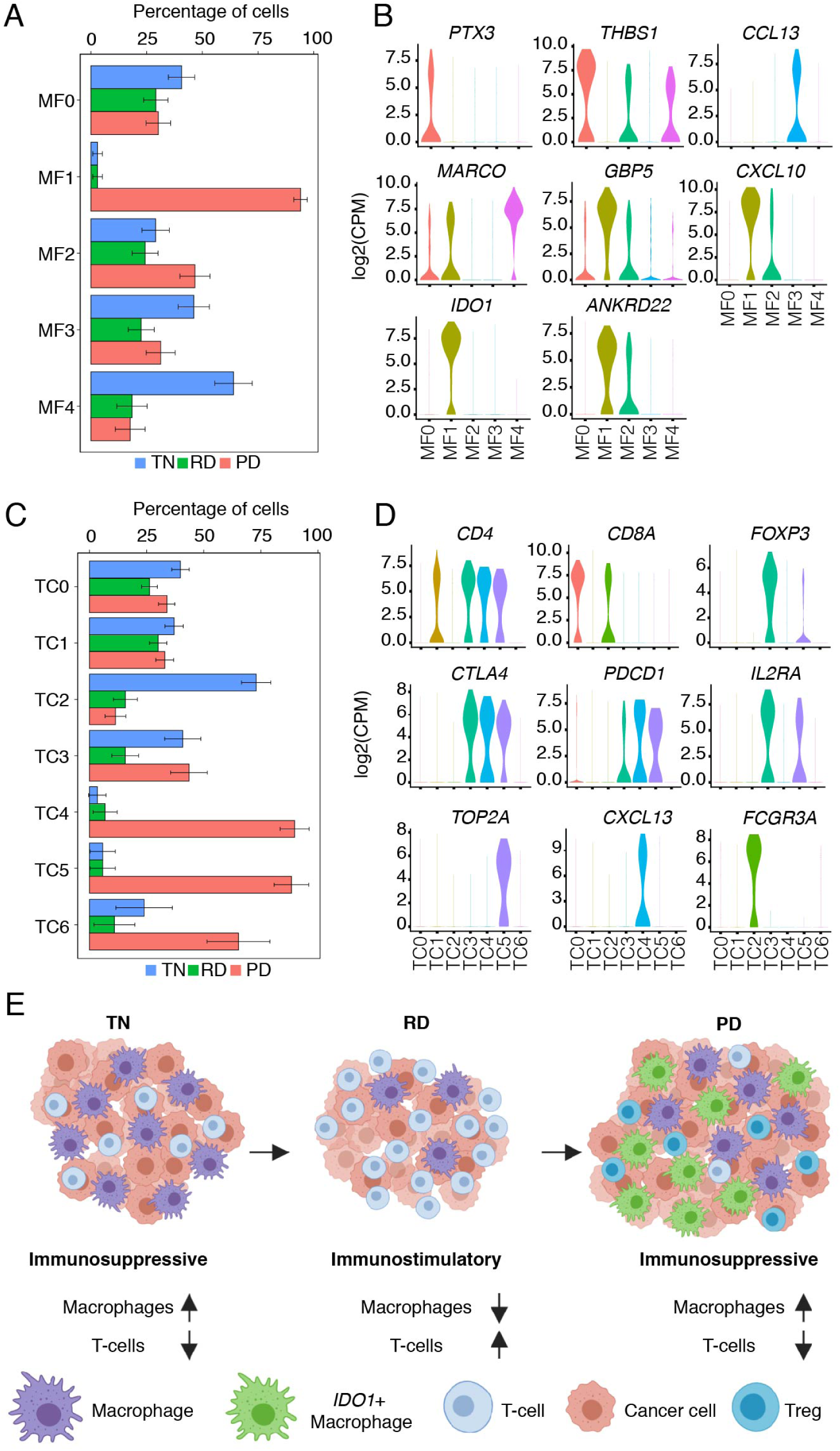
Immune cell subpopulations demonstrate unique transcriptional profiles within each treatment timepoint. (A) Fraction of cells belonging to each treatment stage for each lung-derived macrophage cluster in Supplementary Figure 6. Error bars indicate the 95% confidence interval for the calculated relative frequencies. (B) Violin plots showing the expression level distribution of notable individual genes (C) Fraction of cells belonging to each treatment stage for each lung-derived T-cell cluster in Supplementary Figure 6. Error bars indicate the 95% confidence interval for the calculated relative frequencies. (D) Violin plots showing the expression level distribution of notable individual genes. (E) Graphical summary of immune microenvironment changes across treatment timepoints.

In 2 patients, we examined the immune cells from matched tumor biopsies obtained at different treatment timepoints (TH226 and TH266, Supplemental Figure 5B and C, respectively). In the 2 tumor biopsies available for patient TH266, both macrophages and T-cells showed identical patterns to those observed across the entire cohort: a reduction in the fraction of macrophages and an increase in the fraction of T-cells from TN to RD (Supplemental Figure 5D). TH226 exhibited a similar pattern with the fraction of macrophages decreasing at RD after initiation of treatment and increasing again at PD (Supplemental Figure 5E). In an orthogonal dataset, we performed immunofluorescence analysis using macrophage and T-cell specific markers (see methods, Figure 4D, Supplemental Figure 5F). These analyses validated our scRNAseq findings and demonstrated consistent macrophage and T-cell populations changes during treatment (Figure 4E-F). Additionally, we deconvoluted TCGA bulk transcriptome data for NSCLC into fractions of immune cells types (see Methods) and found that TCGA samples with high fractions of macrophages had significantly worse OS (p<0.01) (Supplemental Figure 5G). This suggests additional clinical relevance of our observations.

These findings are particularly intriguing given their similarity to melanoma tumors treated with PD-1 inhibitor (Riaz et al., 2017), albeit here in the distinct context of oncoprotein-targeted therapy in lung cancer. Specifically, an increase in the number of CD8+ T-cells and NK cells and a decrease in M1 macrophages were observed in melanoma during PD-1 inhibition. There may be common responses in NK/T cells and macrophages during treatment across different tumor histologies and treatments. Hence, conserved approaches to targeting RD across different cancer subtypes and therapeutic modalities may exist, an area for future investigation.

### An *IDO1*-expressing macrophage population is enriched at PD

Macrophages from lung tumor biopsies (n=1,034) were subset and clustered into 5 distinct groups (Supplemental Figure 6A). We then evaluated the differential gene expression in each resulting cluster (Supplemental Figure 6B, Supplemental Table 4). We calculated the fraction of cells originating from each of the three treatment groups in each of the 5 macrophage clusters (Figure 5A).

**Fig. 6.**
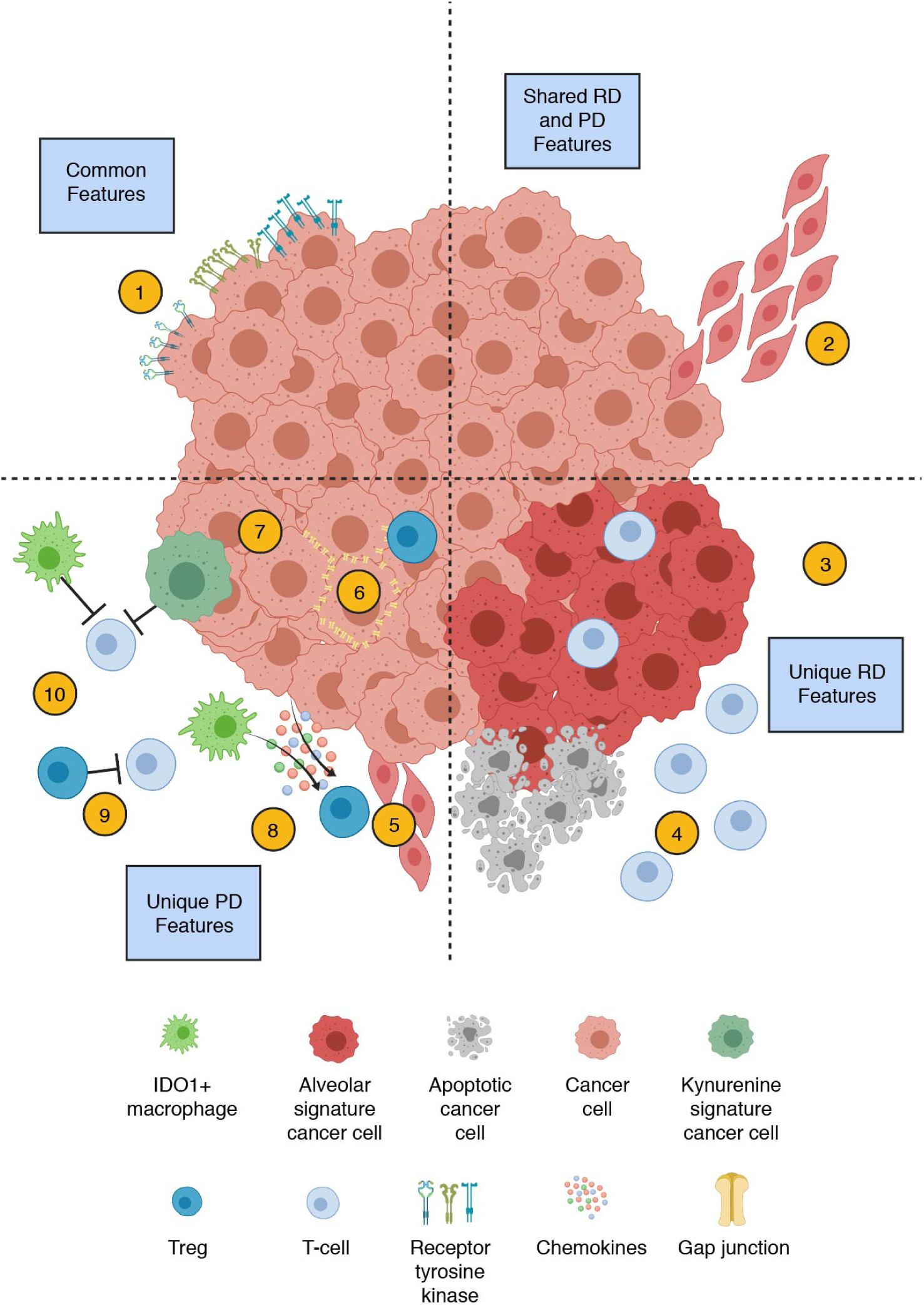
scRNAseq profiles reveal clinical-state specific features of the tumor cellular ecosystem. Features of cancer progression and survival. Common features, found at all treatment timepoints, are shown in the upper left quadrant and include the presence of multiple oncogenic drivers (1). Features shared in RD and PD are shown in the upper right quadrant and include various invasive signaling pathways (2). Features unique at RD, shown in the lower right quadrant, include the Alveolar signature (3) and increased T-cell fraction (4). Features unique to PD, shown in the lower left quadrant, include upregulation of the plasminogen activation pathway (5), expression of gap junction proteins (6), loss of tumor suppressor genes (7), expression of pro-inflammatory chemokines (8), increased Treg population (9), and increased kynurenine signature expression (10).

Cluster MF0, found equally among all three treatment time-points, was enriched for *THBS1* and *PTX3* (Figure 5B, Supplemental Figure 6B), which are associated with resolution of inflammation, wound healing, and with inhibition of IL-1β (Bouhlel et al., 2007; Faz-López et al., 2016; Martinez and Gordon, 2014; Puig-Kröger et al., 2009; Shiraki et al., 2016; Stein et al., 2016). Macrophage clusters MF3 and MF4 were relatively enriched for TN cells. Both clusters were characterized by expression of different genes associated with an immunosuppressive phenotype (*FOLR2, MARCO, RETN, PPARG, CCL13*) (Figure 6B, Supplemental Figure 6B, Supplemental Table 4). Macrophages at PD are overrepresented in group MF1 (Figure 5A), and expressed pro-inflammatory cytokines *CXCL9, CXCL10*, and *CXCL11* (Figure 5B, Supplemental Figure 6B), which favor lymphocyte recruitment into the TME (Nagarsheth et al., 2017). Top differentially expressed genes in this population also included the guanylate-binding family proteins *GBP1*and *GBP5*, which are induced in IFN-γ-activated macrophages and promote inflammatory signaling within the innate immune system via inflamma-some assembly (Shenoy et al., 2012). (Figure 5B, Supplemental Figure 6B).

Despite the expression of pro-inflammatory genes within the MF1 macrophages, the top differentially expressed gene within this group of PD-specific macrophages was *IDO1* (Figure 5B). *IDO1* is induced by inflammation within the TME and promotes a tolerogenic environment through immunosuppressive myeloid cell populations, regulatory T cell (Treg) differentiation, and an immunosuppressive cytokine milieu (Munn and Mellor, 2016).

### An immunosuppressive T-cell phenotype is predominant within the TME at PD

T-cells and NK cells (n=1,725) were analyzed in the same manner as macrophages and resulted in 7 distinct T/NK cell populations (Figure 5C). These included one population (TC2) enriched in TN samples, and 3 populations (TC4, TC5, and TC6) enriched at PD (Figure 5C-D, Supplemental Figure 6D). There was a high fraction of T-cells in RD tumors (Figure 4C) and there was no single T-cell cluster which demonstrated an excess of T-cells in RD (Figure 5C).

Both TN and PD T-cells demonstrated a relative decrease in T-cell infiltration (Figure 4C). TC2 cluster cells, enriched in TN, expressed markers consistent with an NKT cell phenotype, including both T-cell (*CD3, CD8*) and NK cell markers (*KIR2DL3, FCGR3A*), which were less represented in RD and PD samples (Figure 5D, Supplemental Figure 6D). While overall T-cell infiltrate remained limited at PD (Figure 4C), there was relative enrichment for T-cell phenotypes with immunosuppressive features, including T-cell clusters TC4, TC5, and TC6 (Figure 5C). TC4 was identified as a CD4+ T-cell cluster with an exhausted phenotype, characterized by expression of the inhibitory receptors *PDCD1* and *CTLA4* (Wherry and Kurachi, 2015) (Figure 5D). The T-cells within this cluster also expressed *CXCL13*, which suggests that these cells may have features of PD-1-expressing T-follicular helper cells (Tfh). Tfh cells function in crosstalk with the humoral immune system to promote B-cell activation and immunologic memory (Crotty, 2014). TC5 was composed of proliferating Treg cells (expressing *FOXP3, IL2RA, Ki67, TOP2A*). Finally, TC6 is composed of NK cells (*KIR2DL4, GNLY*), the majority of which did not highly express CD3 or *FCGR3A* (CD16) (Figure 5D). This suggests limited cytotoxic function and is consistent with reports of infiltration by poorly cytotoxic NK cells within NSCLC more generally (Carrega et al., 2008), albeit without a prior link to tumors progressing on targeted therapy.

Tumor biopsies obtained at RD revealed the presence of a more pro-inflammatory, “hot”, TME which was absent in TN or PD biopsy samples as manifested by increased overall proportion of T-cells and enrichment of effector T-cell phenotypes (Figure 5E). The majority of the RD T-cells were contained in TC0 and TC1. Cluster TC0 was composed of cytotoxic CD8+ T-cells (*CD8, IFNG, GZMK*), while cluster TC1 was composed of CD4+ T-cells that are negative for co-inhibitory receptor expression (*CD4, CCR7, IL7R*) (Figure 5D, Supplemental Figure 6D), in contrast to the *PDCD1/CTLA4* expressing CD4+ T-cells seen in the PD samples.

In summary, both the TN and PD TME were characterized by the relative predominance of macrophage infiltration over T-cell infiltration; however, the phenotypic characteristics of these infiltrating immune cells differ between the two groups. At PD, there was infiltration by an *IDO1*+ macrophage population, of proliferating regulatory T-cells, and of exhausted CD4+ T-cells which were minimally present at earlier phases of treatment. In contrast, the TN state was characterized by a predominance of more classically immunosuppressive M2-like macrophages (Sica et al., 2008). By distinction, in RD there was increased infiltration of effector T-cell populations with signatures of activity and decreased immunosuppressive macrophage infiltration (Figure 5E).

## Discussion

Though emerging, there remains an incomplete catalogue of single-cell transcriptional data to understand the cell states and therapy-induced evolution of biological heterogeneity of major diseases such as cancer. Precision medicine treatments such as small molecule targeted therapies and immunotherapies have improved cancer patient survival. However, tumors continuously evolve and resistance to these therapies is nearly inevitable. Tumor heterogeneity impacts treatment response and resistance and pertains to cancer cells and the immune and stromal components of the TME (Andor et al., 2016; Morris et al., 2016; Rybinski and Yun, 2016; Swanton, 2012). Cellular plasticity, as a contributor to heterogeneity, has been under-explored in longitudinal clinical tumor biopsies of advanced-stage solid malignancies and is likely to be a key contributor to tumor resilience to systemic therapy and the development of treatment resistance. Single-cell transcriptional data are invaluable in our attempt to understand the cell states and therapy-induced evolution of biological heterogeneity in cancer, particularly in metastatic disease.

Our scRNAseq analyses of advanced-stage NSCLC biopsies obtained longitudinally from individual patients and the clinical cohort as a whole elucidate the rich mutational and transcriptional diversity within individual tumor samples and the dynamic changes in the transcriptional profiles of cancer cells and the TME composition during treatment. These features are clinically-relevant in NSCLC, as we link them to distinct treatment timepoints and clinical states and to differential survival outcomes in independent clinical cohorts such as the MSK-IMPACT and TCGA datasets. Our findings provide a roadmap that highlights the underlying cellular ecosystem and mechanisms to further explore to improve response to treatment.

An important first step was to develop and optimize a clinical pipeline to capture tumor biopsy samples from a treatment phase that is infrequently captured in solid malignancies: namely the phase of RD during initial molecular treatment. Our study offers a rare view of the clinically-relevant biological processes that characterize this poorly-understood phase of the evolution of advanced-stage solid malignancies (here, lung cancer) and creates a resource for both discovery purposes and validation of pre-clinical studies (Blakely et al., 2015; Ramirez et al., 2016; Sharma et al., 2010; Spitzer et al., 2017). We demonstrate successful use of these tumor samples in combination with plate-based scRNAseq in metastatic NSCLC samples. This is critical, as patients with metastatic disease do not routinely receive surgical resection as part of their treatment. Thus, techniques for single-cell profiling that require larger amounts of tissue are not suitable for the interrogation of tissue samples from metastatic disease (Lambrechts et al., 2018; Schelker et al., 2017).

Our scRNAseq data was leveraged to query for the presence of cancer-relevant mutations within expressed regions of the genome. This revealed widespread intra-tumoral heterogeneity in oncogenic alterations that are expressed in cancer cells.

The scRNAseq data complement and extend current clinical analyses which typically: (1) profile a limited subset of genes, (2) provide a view of DNA changes without validation of gene expression, and (3) are performed at bulk cellular scale (Figure 2). Our data suggest that individual cancer cell populations can show expression of the putative oncogenic driver and multiple additional mutations in genes of known oncogenic importance (Figure 6, #1). This is relevant as it provides a potential explanation for why complete responses to treatment are rare. Cancer cells, and their occult genetic subpopulations as revealed by high-resolution scRNAseq, in human tumors already harbor the appropriate genetic framework and evolutionary playbook to evolve resistance. These ‘hard-wired’ properties that can remain undetected by current bulk sampling analysis are further bolstered by the therapy-induced transcriptional plasticity that we demonstrated by longitudinal scRNAseq profiling.

We uncovered transcriptional signatures specific to different treatment time points and clinical states (Figure 3, Figure 6 #3, #5, #6, #7). The majority of these signatures (kynurenine pathway, plasminogen activation pathway, *SERPINE1*, and gap junction genes) were biomarkers of significantly worse overall survival in TCGA lung adenocarcinoma samples and were most pronounced at PD. Conversely, we found the alveolar cell signature was enriched at RD and was associated with improved survival. Our data highlight a connection from the alveolar cell signature to the WNT/ β-catenin pathway as a mechanism of injury-response regeneration. Though the WNT/ β-catenin pathway is potentially therapeutically-targetable, (Krishnamurthy and Kurzrock, 2018) it will be critical to determine how to best modulate this pathway to impact residual cancer cell survival. A general principle our data highlight is that by employing targeted treatments that take advantage of specific cell states we may be able to engineer cancer cell fate(s) to improve therapeutic responses in metastatic solid malignancies, expanding upon this concept in practice in acute promyelocytic leukemia (Chomienne et al., 1990). If deployed at the appropriate time, treatments that target liabilities of a specific cell state or prevent further adaptation may help improve patient survival by constraining continued tumor evolution towards complete drug resistance. It is possible that inhibition of the factors driving the primitive cell state transition at RD could remove this protective biological state and eliminate RD cancer cells. It remains unclear whether the cell state(s) present at RD is unique to cancer cells receiving targeted therapy or also applies to chemotherapy and immunotherapy, an area for future investigation. We have highlighted ongoing efforts targeting signatures we discovered in Table 1 that could be explored to improve targeted therapy response.

Beyond cancer cell-intrinsic signatures, we also explored changes in the TME during treatment. We found a relatively low T-cell infiltration in the TME of TN and PD patients (Figure 5C-D), consistent with prior reports of low cytotoxic T-cell infiltration in treatment-naive EGFR-mutant NSCLC (Gainor et al., 2016). Our results uncovered an induction of a more inflammatory phenotype during RD on targeted therapy, hallmarked by infiltration of cytotoxic T-cells (Figure 6, #4) and decreased infiltration of immunosuppressive macrophages (Figure 5E). This inflammatory state may represent a complement to the alveolar cell, injury-repair and regenerative state present in the cancer cell compartment (described above), with the potential for crosstalk between the cancer cells and TME. These TME changes were transient, as at PD there was enrichment for IDO1-expressing macrophages, regulatory T-cells, and other immunosuppressive T-cell populations. These are all features of an environment hostile to the establishment of an effective immunologic response (Figure 6, #9, #10). The induction of a more immunostimulatory phenotype during targeted therapy (i.e. in RD) may offer a window-of-opportunity to introduce novel TME target-based combination therapies around the time of RD in the context of a more favorable TME to increase initial response and consolidate treatment in a multi-modal approach.

Cancer cell signaling and the TME are linked and there may be treatment strategies which target both compartments concurrently. We identified two such examples: the kynurenine pathway and *SERPINE1*. In the kynurenine pathway, we identified increased pathway activation in cancer cells and myeloid cells at PD (Figure 6, #5). *IDO1*, as a rate limiting enzyme in the kynurenine pathway, can influence diverse components of the TME including T-cell and myeloid cell populations as well as angiogenesis in favor of immunosuppression (Munn and Mellor, 2016). The use of IDO1 inhibitors as part of a combination immunotherapy strategy with PD1/PDL1 checkpoint inhibitors showed promise in early-phase studies (Siefker-Radtke et al., 2018), yet ultimately failed to demonstrate improved outcomes in advanced-stage melanoma (Long et al., 2018). We demonstrated distinct evolving TME states, suggesting that there may be a window-of-opportunity at which point kynurenine pathway inhibitors may be more effective (Figure 6, Table 1). Similarly, SERPINE1 is notable for its activity in both cancer cells, as we identified, and the immune system (Placencio and DeClerck, 2015). Utilizing therapies that leverage this multi-cellular crosstalk may improve patient outcomes.

In summary, the scRNAseq dataset presented here demonstrates the feasibility of performing scRNAseq on tumor biopsies obtained longitudinally at clinically-relevant time-points during the active targeted treatment of advanced-stage solid malignancy patients. Understanding each of the features we discovered and highlighted (Figure 6) in individual cells, as revealed by scRNAseq, is critical to deconvolute tumor biological heterogeneity and evolution during systemic therapy. The data provide a more granular biological foundation to develop strategies for the elimination or neutralization of RD to induce more durable responses for patients with advanced-stage NSCLC and potentially other solid malignancies across different therapeutic modalities.

## ACKNOWLEDGEMENTS

This project is supported by the NIH / NCI U54CA224081, R01CA204302, R01CA211052 and R01CA169338 (to T.G.B.), the Van Auken Foundation (to T.G.B. and C.M.B.), and Novartis Pharmaceuticals (to T.G.B), AstraZeneca, as wells grants from the University of California Cancer League (to C.E.M), The Damon Runyon Cancer Research Foundation P0528804 (to C.M.B), Doris Duke Charitable Foundation grant # 2018110 (to C.M.B), and V Foundation V2018-020 (to C.M.B.), K12 CA086913 (to E.S). F.H. was supported by the Mildred Scheel postdoctoral fellowship from the German Cancer Aid. E.A.Y is supported by T32 HL007185 from the NHLBI. E.L.S is supported by K12 CA086913. Special thanks to Bing Wu and Lillian Cohn for their insights and support.

## Author Contributions

Conceptualization, T.G.B, C.M.B., R.C.D., J.W. S.D. C.E.M., A.M.; Methodology, T.G.B, C.M.B., R.C.D., S.D. C.E.M., A.M., J.K.R.; Software Programming, A.M., L.H. S.D., Wei.W.; Validation, F.H., D.L.K., C.E.M, A.M., P.G., E.L.S., E.A.Y., J.K.R.; Formal Analysis, A.M., L.H. S.D., A.Z., W.T., M.T., R.S., K.A.Y., C.E.M., W.W, J.K.R, E.A.Y, D.L.K, F.H,; Resources, C.E.M. C.M.B, J.K.R., A.M., S.D., N.N., T.L., A.U., K.J, P.K.K., E.S., Y.G., D.M.N., N.J.T., A.G., M.Go, H.D., L.T, M.Gu., T.J. J.R.K., D.J., E.L.S., E.A.Y.; Data Curation Management, A.M, C.E.M.,; S.D.; Writing-Original Draft C.E.M., A.M., J.K.R., S.D.; Writing – Revisions and Editing, All Authors; Visualization, A.M., C.E.M., J.K.R. S.D., C.M.B., T.G.B.; Supervision, S.D., C.M.B., T.G.B.; J.W.; R.C.D., R.G., N.N.; Project Administration, S.D., C.M.B., T.G.B; Funding Acquisition, S.D., C.M.B., T.G.B, C.E.M.

## Code

All code can be found on github 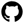czbiohub/scell_lung_adenocarcinoma and 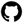czbiohub/cerebra.

## Materials and Methods

### Patient population

All patients gave informed consent for collection of clinical correlates, tissue collection, research testing under Institutional Review Board (IRB)-approved protocols (CC13-6512 and CC17-658, NCT03433469). Formalin-fixed paraffin embedded (FFPE), frozen, and fresh tissue samples were obtained according to the safety standards of the interventional radiologist, pulmonologist, or surgeon. Demographic and clinical history for each patient was obtained from chart review. Days until progression were determined based on imaging studies which demonstrated definitive growth of a known tumor site or new extra-CNS metastatic deposits. Residual disease state was determined by serial imaging demonstrating continued reduction or stability tumor with no evidence of progression. Patient studies were conducted according to the Declaration of Helsinki, the Belmont Report, and the U.S. Common Rule.

### Sample preparation of core and resection samples

Tissue was first cut sample into small pieces and placed into a 1.5 mL tube (or multiple tubes if necessary). 1.5 mL of collagenase buffer (10mL DMEM (GE Life Sciences, SH30081.01), 0.20 g Collagenase Type 2 (Worthington Biochemical, LS004176)) was added to the tube and the sample was digested for 30 minutes at 37°C, shaking in a thermomixer @ 800-1000 rpm. The sample was manually agitated by pipetting up and down 15 times then returned to the thermomixer for 25 minutes. After incubation, the sample was removed from the thermomixer, agitated again by pipetting the sample up and down 15 times before passing the sample through a 100-micron filter (Fisherbrand, 22363548) into a new 15 mL falcon tube. The filter was washed with 1-2 mL of collagenase buffer before the sample was spun in the centrifuge at 500xg for 10 minutes. If the resulting cell pellet was red, 0.5 mL RBC lysis buffer (Thermo Fisher Scientific, A1049201) was added to sample tubes and allowed to sit at room temperature for 3 minutes before quenching with 1.0 mL DMEM (GE Life Sciences, SH30081.01) + 6% FBS (Omega Scientific, Inc, FB-11) and spun in the centrifuge at 500xg for 5 minutes. Remaining cells were stained with 10 µl CD45-FITC (Miltenyi Biotec, 130-080-202) and 1 µl of Hoechst stain (Thermo Fisher Scientific, H3570). Samples incubated on ice in the dark for 20 minutes. One mL of FACS Buffer was then added to the stained cells and spun at 500xg for 10 minutes before aspirating off supernatant. Cells were resuspended with 0.5 mL of FACS Buffer. PI (Life Technologies, P3566) was added immediately prior to sorting (protocols.io: dx.doi.org/10.17504/protocols.io.65rhg56).

### Sample preparation of thoracentesis

samples Cells were filtered through a 100 µm strainer (Fisherbrand 22363548), pelleted (500xg, 5 min, 4°C), and resuspended in FACS buffer. Cells were then stained with CD45-FITC (Miltenyi Biotec, 130-080-202) for 20 min at 4°C in the dark. Cells were then pelleted (500xg, 5 min, 4°C) and resuspended in FACS buffer before being transferred to a FACS tube (Falcon 14-956-3C). Sytox Blue dead cell stain (Thermo Fisher Scientific, S34867) was added immediately prior to sorting.

### Lysis plate preparation

Lysis plates were created by dispensing 0.4 µl lysis buffer (0.5U Recombinant RNase Inhibitor (Takara Bio, 2313B), 0.0625% TritonTM X-100 (Sigma, 93443-100ML), 3.125 mM dNTP mix (Thermo Fisher, R0193), 3.125 µM Oligo-dT30VN (IDT, 5’AAGCAGTGGTATCAACGCAGAGTACT30VN-3’) and 1:600,000 ERCC RNA spike-in mix (Thermo Fisher, 4456740)) into 384-well hard-shell PCR plates (Biorad HSP3901) using a Tempest liquid handler (Formulatrix). All plates were then spun down for 1 minute at 3220xg and snap frozen on dry ice. Plates were stored at -80°C until used for sorting.

### FACS sorting

Cells were sorted into 384-well plates using SH800S (Sony) sorter. Cells were sorted using the “Ultra purity” setting on the sorter. For a typical sort, a tube containing 0.3-1ml the pre-stained cell suspension was vortexed gently and loaded onto the FACS machine. A small number of cells were flowed at low pressure to check cell concentration and amount of debris. Then the pressure was adjusted, flow was paused, the first destination plate was unsealed and loaded. Single-cell sorting was done where half the plate was sorted for CD45+/PI-/Hoechst+ while the second half was sorted for CD45-/PI-/Hoechst+. Immediately after sorting, plates were sealed with a pre-labeled aluminum seal, centrifuged and flash frozen on dry ice.

### cDNA synthesis and library preparation

cDNA synthesis was performed using the Smart-seq2 protocol (Picelli et al., 2013, 2014). Briefly, 384-well plates containing single-cell lysates were thawed on ice followed by first strand synthesis. 0.6 µl of reaction mix (16.7 U/µl SMARTScribe Reverse Transcriptase (Takara Bio, 639538), 1.67 U/µl Recombinant RNase Inhibitor (Takara Bio, 2313B), 1.67X First-Strand Buffer (Takara Bio, 639538), 1.67 µM TSO (Exiqon, 5’-AAGCAGTGGTATCAACGCAGACTACATrGrG+G-3’), 8.33 mM DTT (Bioworld, 40420001-1), 1.67 M Betaine (Sigma, B0300-5VL), and 10 mM MgCl2 (Sigma, M1028-10X1ML)) was added to each well using a Tempest liquid handler or Mosquito (TTP Labtech). Reverse transcription was carried out by incubating wells on a ProFlex 2×384 thermal-cycler (Thermo Fisher) at 42°C for 90 min and stopped by heating at 70°C for 5 min.

Subsequently, 1.5 µl of PCR mix (1.67X KAPA HiFi HotStart ReadyMix (Kapa Biosystems, KK2602), 0.17 µM IS PCR primer (IDT, 5’-AAGCAGTGGTATCAACGCAGAGT-3’), and 0.038 U/µl Lambda Exonuclease (NEB, M0262L)) was added to each well with a Mantis liquid handler (Formulatrix) or Mosquito, and second strand synthesis was performed on a ProFlex 2×384 thermal-cycler by using the following program: 1. 37°C for 30 minutes, 2. 95°C for 3 minutes, 3. 23 cycles of 98°C for 20 seconds, 67°C for 15 seconds, and 72°C for 4 minutes, and 4. 72°C for 5 minutes.

The amplified product was diluted with a ratio of 1 part cDNA to 10 parts 10mM Tris-HCl (Thermo Fisher, 15568025). 0.6 µl of diluted product was transferred to a new 384-well plate using the Viaflow 384 channel pipette (Integra). Illumina sequencing libraries were prepared as described in (Darmanis et al., 2015). Briefly, tagmentation was carried out on double-stranded cDNA using the Nextera XT Library Sample Preparation kit (Illumina, FC-131-1096). Each well was mixed with 0.8 µl Nextera tagmentation DNA buffer (Illumina) and 0.4 µl Tn5 enzyme (Illumina), then incubated at 55°C for 10 min. The reaction was stopped by adding 0.4 µl “Neutralize Tagment Buffer” (Illumina) and spinning at room temperature in a centrifuge at 3220xg for 5 min. Indexing PCR reactions were performed by adding 0.4 µl of 5 µM i5 indexing primer, 0.4 µl of 5 µM i7 indexing primer, and 1.2 µl of Nextera NPM mix (Illumina). All reagents were dispensed with the Mantis or Mosquito liquid handlers. PCR amplification was carried out on a ProFlex 2×384 thermal cycler using the following program: 1. 72°C for 3 minutes, 2. 95°C for 30 seconds, 3. 12 cycles of 95°C for 10 seconds, 55°C for 30 seconds, and 72°C for 1 minute, and 4. 72°C for 5 minutes.

### Library pooling, quality control, and sequencing

Following library preparation, wells of each library plate were pooled using a Mosquito liquid handler. Pooling was followed by two purifications using 0.7x AMPure beads (Fisher, A63881). Library quality was assessed using capillary electrophoresis on a Fragment Analyzer (Agilent) or Tapestation (Agilent), and libraries were quantified by qPCR (Kapa Biosystems, KK4923) on a CFX96 Touch Real-Time PCR Detection System (Biorad). Plate pools were normalized to 2 nM and equal volumes from library plates were mixed together to make the sequencing sample pool.

### Sequencing libraries from 384-well plates

Libraries were sequenced on the NextSeq or NovaSeq 6000 Sequencing System (Illumina) using 2 x 100bp paired-end reads and 2 x 8bp or 2 x 12bp index reads. NextSeq runs used high output kits, whereas NovaSeq runs used either a 200 or 300-cycle kit (Illumina, 20012860). PhiX control library was spiked in at 1%.

### Alignment and gene counts

Sequences from the Illumina sequencing were demultiplexed using bcl2fastq version 2.19.0.316. Reads were aligned using the hg38 genome using STAR version 2.5.2b with parameters TK. Gene counts were produced using HTSEQ version 0.6.1p1 with default parameters except stranded was set to false and mode was set to intersection-nonempty.

### General clustering

Standard procedures for filtering, variable gene selection, dimensionality reduction, and clustering were performed using the Seurat package (Satija et al., 2019) (version 2.3.4) in R (R Core Team (2013). R: A language and environment for statistical computing. R Foundation for Statistical Computing, Vienna, Austria. http://www.R-project.org/), where cells with fewer than 500 genes and 50,000 reads were excluded. Samples with less than 10 total cells were filtered from the analysis. Counts were log-normalized, then scaled by linear regression against the number of reads. Variable genes (Ngenes=6,112) were selected using a threshold for dispersion, with z-scores normalized by expression level. The variable genes were projected onto a low-dimensional subspace using principal component analysis. The number of principal components (Npcs) were selected based on inspection of the plot of variance explained (Npcs = 20). A shared-nearest-neighbors graph was constructed based with metric the Euclidean distance in the low-dimensional subspace. Cells were visualized using a 2-dimensional tSNE on the same distance metric (Res = 0.5, Kparam = 30, script 03). Cell types were assigned to each cluster of cells using the abundance of known marker genes (Supplemental Table 2, script S01-03 and script NI01).

### Epithelial subset clustering, identification of tumor cells, and annotation of non-tumor epithelial cells

Cells previously annotated as epithelial (n = 5,109) were subset and re-clustered using methods described above and the following parameters: Ngenes = (6,919), Npcs = 20, Res = 0.9, Kparam = 10 (script NI02). Malignant epithelial cells were identified using inferCNV (Tickle T, Tirosh I, Georgescu C, Brown M, Haas B (2019). inferCNV of the Trinity CTAT Project. Klarman Cell Observatory, Broad Institute of MIT and Harvard, Cambridge, MA, USA.https://github.com/broadinstitute/inferCNV.). inferCNV which works by finding cells with large copy number variations as determined by sorting expressed genes by their chromosomal location and applying a moving average, a sliding window of 100 genes within each chromosome, to the relative expression values (Patel et al., 2014; Puram et al., 2017; Tirosh et al., 2016). All epithelial cells as well as 300 fibroblasts and 300 endothelial cells were used as input (script NI03). An additional 500 fibroblasts and 500 endothelial cells were used as reference controls. We scored each cell for the extent of CNV signal and plotted cells on a dendrogram which was then cut at the highest point in which all the spiked in endothelial and fibroblasts cells belonged to one cluster (k = 3). All cells that clustered together with spiked in controls were labeled “nontumor”, whereas the remaining two clusters were labeled as “tumor”.

Nonmalignant epithelial cells (n = 1,489), as determined as those cells lacking large chromosomal aberrations from InferCNV analysis, were subset and re-clustered using the following parameters: Ngenes = (7,213), Npcs = 20, Res = 0.3, Kparam = 30 (script NI05). Cell types were assigned to each cluster of cells using the abundance of known marker genes (Supplemental Table 2) and differentially expressed genes as found by using the Seurat function *FindAllMarkers* using the default Wilcoxon rank sum test (arguments used: only.pos = TRUE, min.pct = 0.25, thresh.use = 0.25).

### Tumor cell subset clustering and differential gene expression

Malignant epithelial cells (n = 3,620), as determined as those cells harboring large chromosomal aberrations from InferCNV analysis, were subset and re-clustered using the following parameters: Ngenes = 6,797, Npcs = 15, Res = 0.9, Kparam = 30 (script NI04). We found the differences in gene expression between the three treatment response groups (TN, RD, and PD) using the Seurat function *FindMarkers* using the default Wilcoxon rank sum test. Three separate tests were used to ascertain the differences between: 1) TN and RD, 2) TN and PD and 3) RD and PD (Supplemental Tables 5). Resulting differential gene lists were then filtered to limit patient specific effects. This is achieved by setting a threshold for non-zero expressing cells per patient (RD = 3, 37% of RD patients and PD = 6, 33% of PD patients) and removing differentially expressed genes explained by less than the thresholds set. The top 100 genes from each comparison were manually curated to evaluate for pathway activation. Decreased expression could indicate lack of detection due to the stochasticity of scRNAseq and thus for analysis of activated pathways we focused on upregulated genes. Gene signatures (Supplemental Table 2) were compiled using differential expressed as well as known cell marker genes. Specifically, the alveolar signature (Supplemental Figure 3B) is made of differentially expressed AT1/AT2 genes among the cancer cell timepoint comparisons as well has additional known AT1/AT2 genes (Vieira Braga et al., 2019; Wade et al., 2006). The remaining signatures were identified directly from top differentially expressed genes.

To ensure that we were not misclassifying healthy AT2 cells as cancer cells, we compared the expression levels of our combined alveolar gene signature between the three timepoints (TN, RD, PD) and non-cancer AT2 cells from our dataset as well as additional non-cancer AT2 cells from an external dataset (Vieira Braga et al., 2019). Non-cancer AT2 cells from our dataset were more similar to the external AT2 cells than any of our cancer cells across all timepoints (ρ=0.73, -0.14, 0.23, -0.19, for non-cancer AT2 cells, and TN, RD, PD cancer cells respectively) (Supplemental Figure 3C).

Longitudinal analysis of a single patient was done by subsetting all cell originating from patient TH226. As above, the differences in gene expression between the three treatment response groups (TN, RD, and PD) was found by applying the Seurat function *FindMarkers* using the default Wilcoxon rank sum test. Three separate tests were used to ascertain the differences between: 1) TN and RD, 2) TN and PD and 3) RD and PD (script NI07-08, Supplemental Table 5)

### Survival analysis using cancer cell gene signatures within the TCGA

TCGA LUAD data were downloaded from xenabrowser.net/datapages/. Metadata was downloaded from An Integrated TCGA Pan-Cancer Clinical Data Resource (Liu et al., 2018a). Mean expression of each cancer cell expression signature (alveolar, kynurenine, plasminogen activating, *SER-PINE1*, and gap junction) was calculated per TCGA sample. TCGA samples were then split by quartile groups. Only quartile 4 (high expression) and quartile 1 (low expression) were plotted using library packages survival (Therneau T (2015). A Package for Survival Analysis in S. version 2.38, https://CRAN.R-project.org/package=survival.) and survminer (Aldoukadel Kassambara, Marcin Kosinski and Przemylslaw Biecek (2019). survminer: Drawing Survival Curves using ‘ggplot2’. R package version 0.4.5. http://CRAN.R-project.org/package=surviminer) in R (script NI10).

### Immunohistochemistry

All specimens were acquired from individuals with NSCLC as noted above. 4-micron thick formalin-fixed paraffin embedded (FFPE) human tissue sections were processed using previously published method (Haderk et al., 2019) and manufacturer’s recommendations for antigen retrieval. Staining was performed overnight AQP4 rabbit monoclonal antibody (#59678, Cell Signaling Technology, 1:100 dilution). Stained slides were digitized using an Aperio ScanScope XT Slide Scanner (Leica Biosystems) with an either an 20X or 40X objective. Tumor populations were annotated, then scored in a blinded, randomized analysis by a pathologist for percent tumor positivity and subcellular staining intensity at the membrane, cytosolic, and nuclear compartments. Staining intensity was graded as negative, weak, intermediate, or strong and received scores of 0, 1, 2, or 3 respectively. Percent tumor positivity coefficient was graded as 0, negative; 1, less than 10% immunopositive; 2, between 10-50% immunopositive; 3, between 51-80% immunopositive; 4, greater than 80% immunopositive. Calculation of immunoreactivity scores was performed by multiplying the staining intensity score (0-3) with the percent tumor positive coefficient (0-4) to yield a value between 0 and 12 (Fedchenko and Reifenrath, 2014).

### Mutation detection from scRNAseq

Alignment bams for all non-immune cells (stroma and epithelial) were passed to GATK HaplotypeCaller which was run from the latest available Docker container (broadinstitute/gatk:4.0.11.0) using the following options:

*–disable-read-filter MappingQualityReadFilter*
*–disable-read-filter GoodCigarReadFilter*
*–disable-read-filter NotSecondaryAlignmentReadFilter*
*–disable-read-filter MappedReadFilter*
*–disable-read-filter MappingQualityAvailableReadFilter*
*–disable-read-filter NonZeroReferenceLengthAlignmentReadFilter*
*–disable-read-filter NotDuplicateReadFilter*
*–disable-read-filter PassesVendorQualityCheckReadFilter*
*–disable-read-filter WellformedReadFilter*

Disabling these specific read filters proved necessary for scRNAseq, as inherent low-coverage causes the vast majority of reads to be flagged for removal otherwise. The full human variant set (dbSNP) was downloaded from NCBI (https://www.ncbi.nlm.nih.gov/variation/docs/human_variation_vcf/), and every variant call was assessed for its presence/absence in the human variant database. dbSNP is a public, living catalog of 674 million human somatic SNPs and indels that have been reported by peer-reviewed publications.

Cloud-based parallelization of HaplotypeCaller jobs was achieved with Reflow, a workflow engine for distributed, incremental data processing in the cloud (https://github.com/grailbio/reflow). HaplotypeCaller outputs a separate variant calling format file (VCF) for each cell, which were processed with the python package cerebra (https://github.com/czbiohub/cerebra). Variants found in dbSNP were removed, not to be included in further analysis. We reasoned that by removing ‘common’, population-level variants, we could better hone-in on disease specific variation.

In addition to scRNAseq reads, we obtained bulk DNA reads from peripheral blood for the majority of our patients (with the exception of three). These PBMC reads were run through HaplotypeCaller to establish ‘germline’ mutation profiles for each of our patients. Germline mutations were then subtracted out from each of that patient’s single cell VCFs. This filtering step was omitted for the three patients for which we did not obtain peripheral blood, however, these single cell VCFs were still passed through our dbSNP filter.

We also applied a *fathmm* filter to all cells (Shihab et al., 2015). *fathmm* takes a machine learning approach to predict the likelihood of a given SNP to be pathogenic, integrating ENCODE annotations for things like transcription factor binding sites, histone modifications, cross-species sequence alignment and conservation scores, etc. Only variants computationally predicted to be pathogenic were included in our analysis, i.e. those variants with a *fathmm* score > 0.7.

The remaining variants were then filtered through the COSMIC (Catalogue of Somatic Mutations in Cancer) complete mutation – genome screens database (https://cancer.sanger.ac.uk/cosmic/download). Only SNPs/indels associated with ‘Lung’ as per their COSMIC annotation were kept. Variant calls were mapped to their corresponding genes, and per-patient / per-sample mutational profiles were established. We used the ERCCs spiked into each cell sample as a negative control for false positive mutations, which can arise due to technical artifacts such as PCR errors. We found the median false positive mutation rate to be 0.000256% per base (Enge et al., 2017).

### Fusion detection from scRNAseq

Fusion transcripts were detected with STAR-fusion (https://github.com/STAR-Fusion/STAR-Fusion/wiki) version 1.6.0, run from a Docker container (trinityctat/ctatfusion:1.5.0). The following options were used:

*–FusionInspector validate*
*–examine_coding_effect*
*–denovo_reconstruct*

Distributed processing of STAR-fusion jobs was accomplished with Reflow. Output files were processed with cerebra, then combined with variant calls to create per-cell and per-sample summary tables.

### Mutational analysis of tumor cells

Information extracted from cerebra was summarized by sample. Coverage information was provided by a secondary output from cerebra summarized by sample and gene. Where all cells are summarized by sample and all *fathmm* filtered ROIs are summarized by corresponding gene (script NI06). Plots were generated using the R pheatmap package(Raivo Kolde (2019). pheatmap. Pretty Heatmaps. R package version 1.0.12. https://CRAN.R-project.org/package=pheatmap).

Survival analysis using number of mutations within the MSK-Impact data MSK-Impact data was downloaded from cBioPortal (Cerami et al., 2012; Gao et al., 2013) and subset to only NSCLC samples MSK-Impact data was subset to only those mutations that were also found in the scRNAseq dataset of mutations (n=141 unique mutations). We stratified MSK-Impact samples by those with greater than or equal to 2 mutations from the tier one COSMIC mutations found in the scRNAseq dataset (mutation high), and those less than 2 mutations (mutation low) (Figure 2D). Kaplan-Meier plots were visualized with the lifelines package in python (Cameron Davidson-Pilon, Jonas Kalderstam, Paul Zivich, Ben Kuhn, Andrew Fiore-Gartland, Luis Moneda, … André F. Rendeiro. (2019, September 4). CamDavidsonPilon/lifelines: v0.22.4 (Version v0.22.4). Zenodo. http://doi.org/10.5281/zenodo.3386382; Python Software Foundation. Python Language Reference, version 3.4. Available at) (script NI12).

### Immune subset clustering and differential gene expression

All cells annotated as immune (n=12,077) were subset and clustered as described above (script IM01) using the following parameters (Ngenes=1,690, Npc=20, Res=0.3, k.param=10). The resulting 17 clusters were assigned to different major immune cells types using a list of curated gene markers (Supplemental Table 2) and by manual curation of differentially expressed genes for each cluster (Supplemental Table 5). The different cell types and number of cells belonging to each type are described in the main text.

To assess changes in fractional abundance of different immune cell populations we used all cells though excluded thoracentesis and brain samples due to difference in the immune makeup of these tumor environments which would skew the data. The function freqCI from the R package REdaS (Maier MJ (2015). Companion Package to the Book R: Einfuhrung durch angewandte Statistik. R package version 0.9.3. htt://CRAN.R-project.org/package=RedaS) (script IM02) was used to calculate confidence intervals for relative frequencies.

Macrophages (n=1,034) and T-cells (n=1,725) from lung biopsies were subset and clustered as described above (script IM03 and IM04 respectively) using the following parameters for MFs (Ngenes=1,637, Npc=10, res=0.3, k.param=10) and T-cells (Ngenes=1,475, Npc=10, res=0.3, k.param=10). The resulting clusters are discussed in the main text and the lists of differentially expressed genes are provided (Supplemental Table 5). We repeated this analysis where we subset the data to only patients with multiple biopsies and sufficient cells (TH226 and TH266) (script IM05).

### Multiplex Immunofluorescence

Multiplex immunofluorescence staining was performed on sequential 4 micron FFPE slides, utilizing the Opal IHC Multiplex Assay (Perkin Elmer). Slides were stained with one of two 6 antibody panels. Panel 1 contained primary antibodies against PD-L1 (clone E1L3N, dilution 1:50, Cell Signaling Technologies), CD68 (clone KP1, dilution 1:500, Dako), IDO (clone D5J4E, dilution 1:100, Cell Signaling Technologies), HLA-DR (clone CR3/43, dilution 1:250, Abcam), CD14 (clone SP192, dilution 1:100, Abcam), and cytokeratin (polyclonal Z0622, dilution 1:250, Dako). Panel 2 consisted of primary antibodies against CD3 (clone LN10, Leica), PD-1 (clone NAT105, dilution 1:100, Abcam), CD14, CD8 (clone C8/144B, dilution 1:100, Dako), FoxP3 (clone 236A/E7, dilution 1:200, Abcam), and cytokeratin. Whole slide scans were acquired at 10X via the Vectra imaging system (Perkin Elmer, version 3.0). Three to six regions from each slide containing tumor and stroma were selected utilizing Phenochart (v1.0.8, Perkin Elmer) for high resolution multispectral acquisition on the Vectra system at 20X magnification. Spectral libraries were generated from single color stains on human tonsil tissue. Spectral unmixing, cell segmentation, and cell phenotype assignment as tumor, macrophage, T cell, or other cell population were performed utilizing InForm image analysis software (Perkin Elmer, version 2.4). Fractions of macrophage and T-cell populations were calculated as: (population of interest) / (macrophage + T-cell populations) and plotted using ‘ggplot2’ in R (H. Wickham. ggplot2: Elegant Graphics for Data Analysis. Springer-Verlag New York, 2016) (script NI15).

### Survival analysis using fractional immune population within the TCGA

As with the survival analysis using cancer cell gene signatures, we used the downloaded TCGA LUAD dataset and metadata to access patient survival outcomes as they pertain to the fractional changes of immune populations within a given tumor. We used CIBER-SORT (Newman et al., 2015) to deconvolute the bulk TCGA samples into relative fractions of immune cell populations as determined by using the LM22 reference. The total macrophage population was found by combining fractions for Monocytes, Macrophages.M0, Macrophages.M1, and Macrophages.M2. The total T-cell population was found by combining fractions of T.cells.CD8, T.cells.CD4.naive, T.cells.CD4.memory.resting, T.cells.CD4.memory.activated, T.cells.follicular.helper, T.cells.regulatory..Tregs, T.cells.gamma.delta, NK.cells.resting, and NK.cells.activated. TCGA samples were then split by quartile groups. Only quartile 4 (high population fraction) and quartile 1 (low population fraction) were plotted using library packages survival (Therneau T (2015). A Package for Survival Analysis in S. version 2.38, https://CRAN.R-project.org/package=survival) and survminer (Aldoukadel Kassambara, Marcin Kosinski and Prze-mylslaw Biecek (2019). survminer: Drawing Survival Curves using ‘ggplot2’. R package version 0.4.5. http://CRAN.R-project.org/package=surviminer) in R (script NI10).

### Generation of in vitro drug tolerant persister and acquired resistance cells and RT PCR

For validation of candidate gene expression via a RT2 Profiler PCR array (Qiagen, CLAH34795), human lung cancer PC9 (EGFR del19) cells were treated for 48 hours (day 2) with DMSO (TN) or for 7 and 19 days with 2µM Osimertinib (Selleck Chemicals LLC, S7297) with replenishment of drug every 3-4 days (RD), respectively. PD samples were derived from an acquired resistant PC9 cell line (Osimertnib IC50 = 89µM), that was generated by continuous treatment with 2µM Osimertinib with replenishment of drug every 3-4 days and presented active proliferation under drug when resistance was called. RNA was extracted via RNeasy Mini Kit (Qiagen, 74104). RNA quality was confirmed as RIN ^3^ 7.5 via Bioanalyzer RNA 6000 Pico kit (Agilent, 5067-1514) and RNA was quantified via Qubit RNA HS Assay kit (Thermo Fisher Scientific, Q32852). A total of 400ng of RNA was reverse transcribed using the First Strand Synthesis Kit (Qiagen, 330401) and then loaded into a custom 384 well RT2 profiler array (Qiagen, CLAH34795). Fold Change was calculated by determining the ratio of mRNA levels to control (day 2) values using the delta threshold cycle (Ct) method (DCt). A t-test was used to find the significance of change between baseline (day 2) and treated timepoints (days 7, 19 and 70) based on normalized Cts to baseline (script NI14).

## Supplementary Figures

**Fig S1 related to Figure 1.**
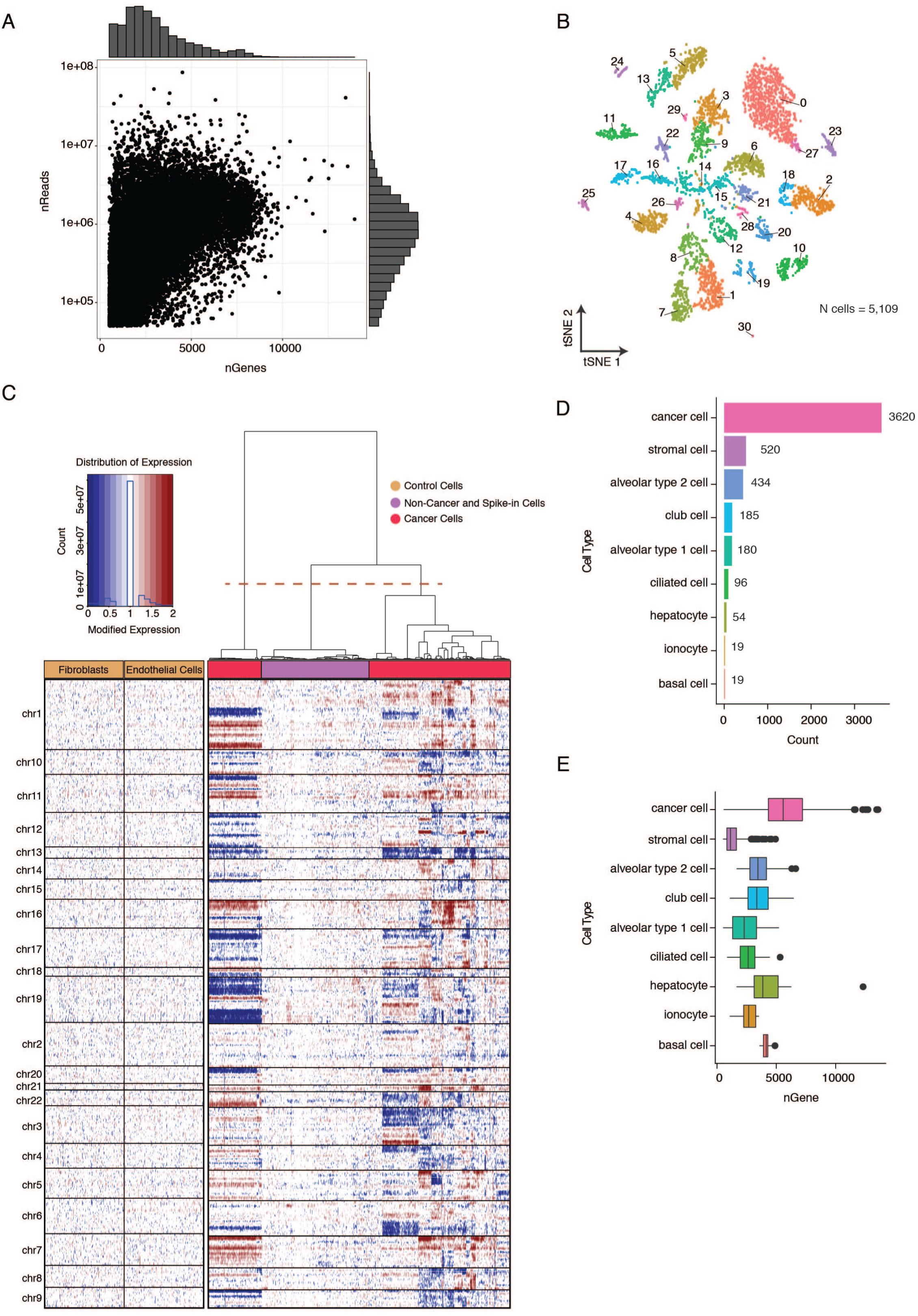
(A) Scatterplot with marginal histograms showing the number of unique genes plotted against the number of reads for all cells passing quality control. (B) t-SNE of all epithelial cells (n=5109), numbers correspond to individual clusters. (C) Inferred large-scale copy number variations (CNVs) help identify cancer (pink) and non-cancer cells (purple). Epithelial and spike in control cells are included in the x-axis and chromosomal regions on the y-axis. Amplifications (red) or deletions (blue) were inferred by averaging expression over 100-gene stretches on the respective chromosomes. (D) Bar plot of cell counts for annotated epithelial cells. (E) Bar plot of the number of unique genes across all annotated epithelial cell types.

**Fig S2 related to Figure 2.**
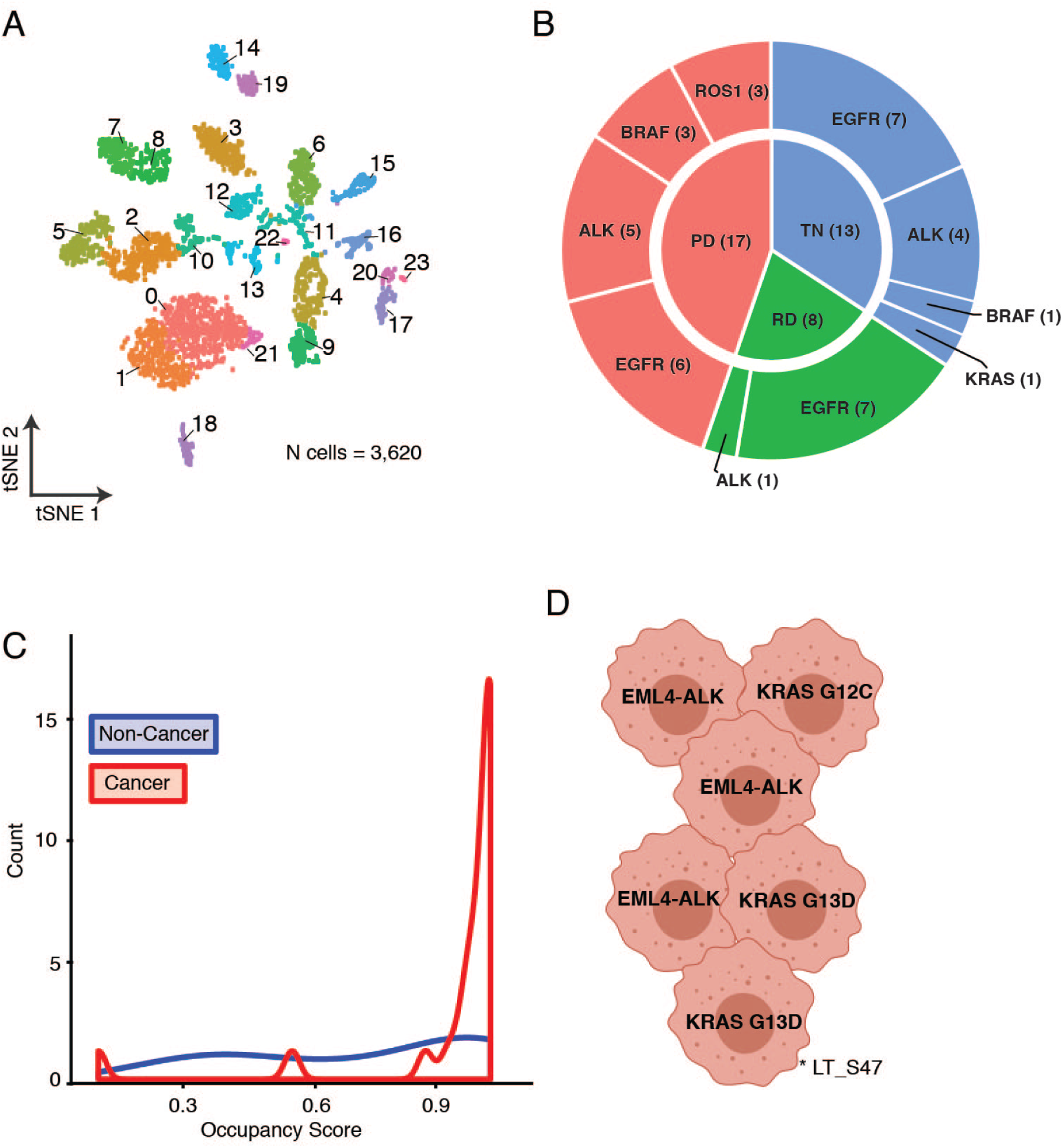
(A) t-SNE plot of 3620 cancer cells from 38 samples, numbers indicate individual clusters. (B) Circle plot illustrating the clinically identified oncogenic driver (outer circle) and timepoints (inner circle) of each biopsy, only for cancer cells. C) Density distribution of cluster occupancy of cancer (red) and non-cancer (blue) epithelial cell clusters, calculated as the percentage of the highest contributing individual patient over the total number of cells for that cluster. (D) Illustration of heterogeneity of primary driver mutated cancer cells found in exemplary sample LTS47 (* in Figure 2A).

**Fig S3 related to Figure 3.**
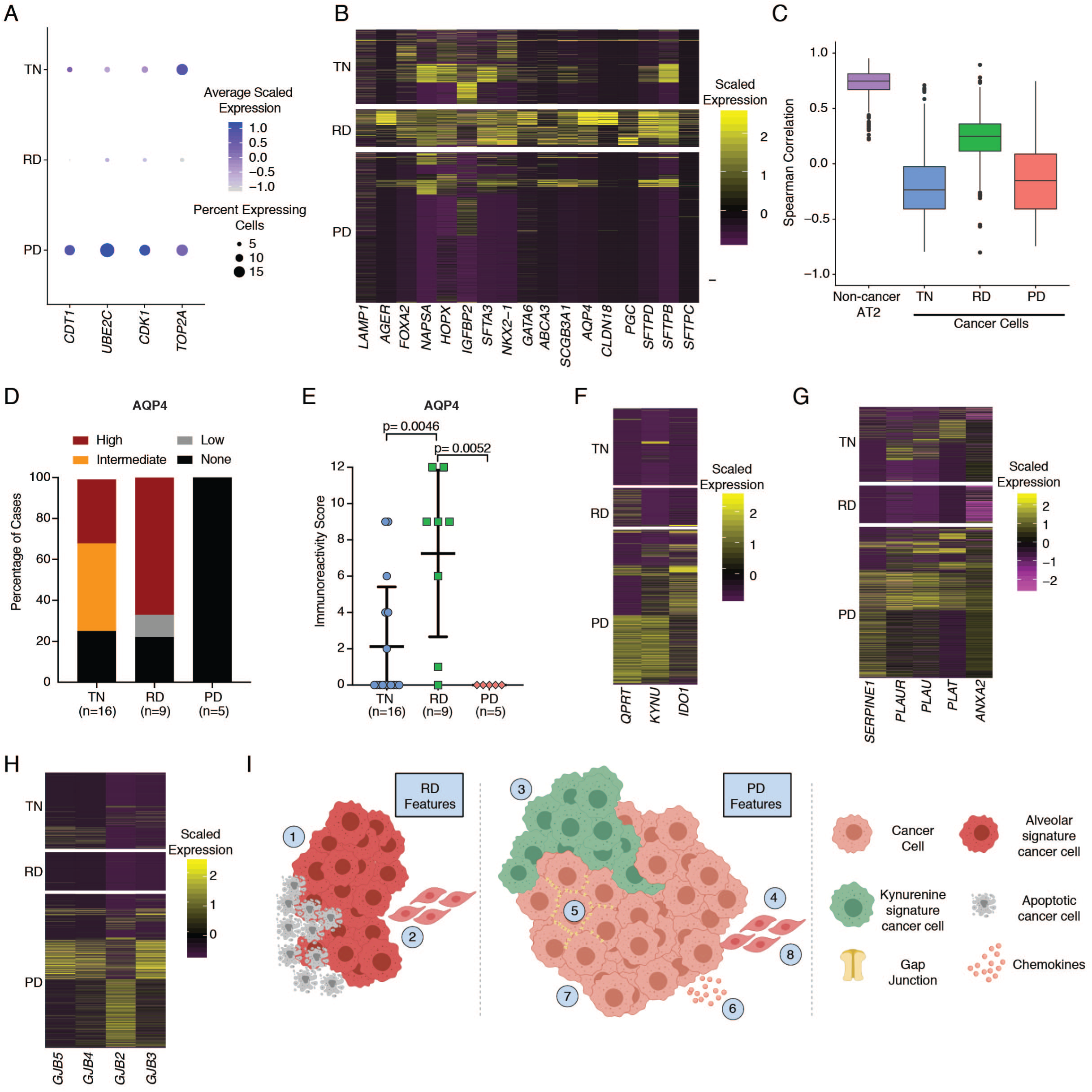
(A) Dot plot of the relative expression of established cellular proliferation genes (x-axis) across treatment timepoints (y-axis). The color intensity scale reflects the average gene expression and the size scale indicates the number of cells expressing the gene within that treatment timepoint. Applying grouped, pairwise comparisons of treatment timepoints of the average scaled expression of all genes demonstrated significantly different expression (p < 0.000001) in all comparisons. (B) Heatmap showing the expression of genes in the alveolar signature. Cells are grouped by treatment timepoint. (C) Box plot of Spearman correlations of cancer cells from all treatment timepoints and healthy AT2 cells to an external reference of healthy AT2 cells. Non-cancer AT2 cells from our dataset were more similar to the external, healthy AT2 cells than any of our cancer cells across all timepoints (mean spearman correlation=0.73, -0.14, 0.23, -0.19, for healthy AT2 cells, and TN, RD, PD cancer cells, respectively. (D) Scoring of IHC stains for AQP4 in TN (n=19), RD (n=9), and PD (n=5) tumor tissue sections. Chi-square testing demonstrated a significant difference (p = 0.0038) in proportions of staining intensity groups between treatment timepoints. (E) Immunoreactivity score for AQP4 across all timepoints from S3D. (F, G, H) Heatmaps showing the expression of genes within each signature (kynurenine, SERPINE1/plasminogen activation, and gap junction, respectively) grouped by treatment timepoint. (I) Graphical summary of cancer cell expression changes across treatment timepoints. RD features include (1) Alveolar signature, and (2) various RD specific invasive signaling pathways. PD features include: (3) kynurenine signature, (4) plasminogen activation and SERPINE1 signatures, (5) gap junction proteins, (6) expression of pro-inflammatory chemokines, (7) loss of tumor suppressor genes, and (8) various PD specific invasive signaling pathways.

**Fig S4 related to Figure 3.**
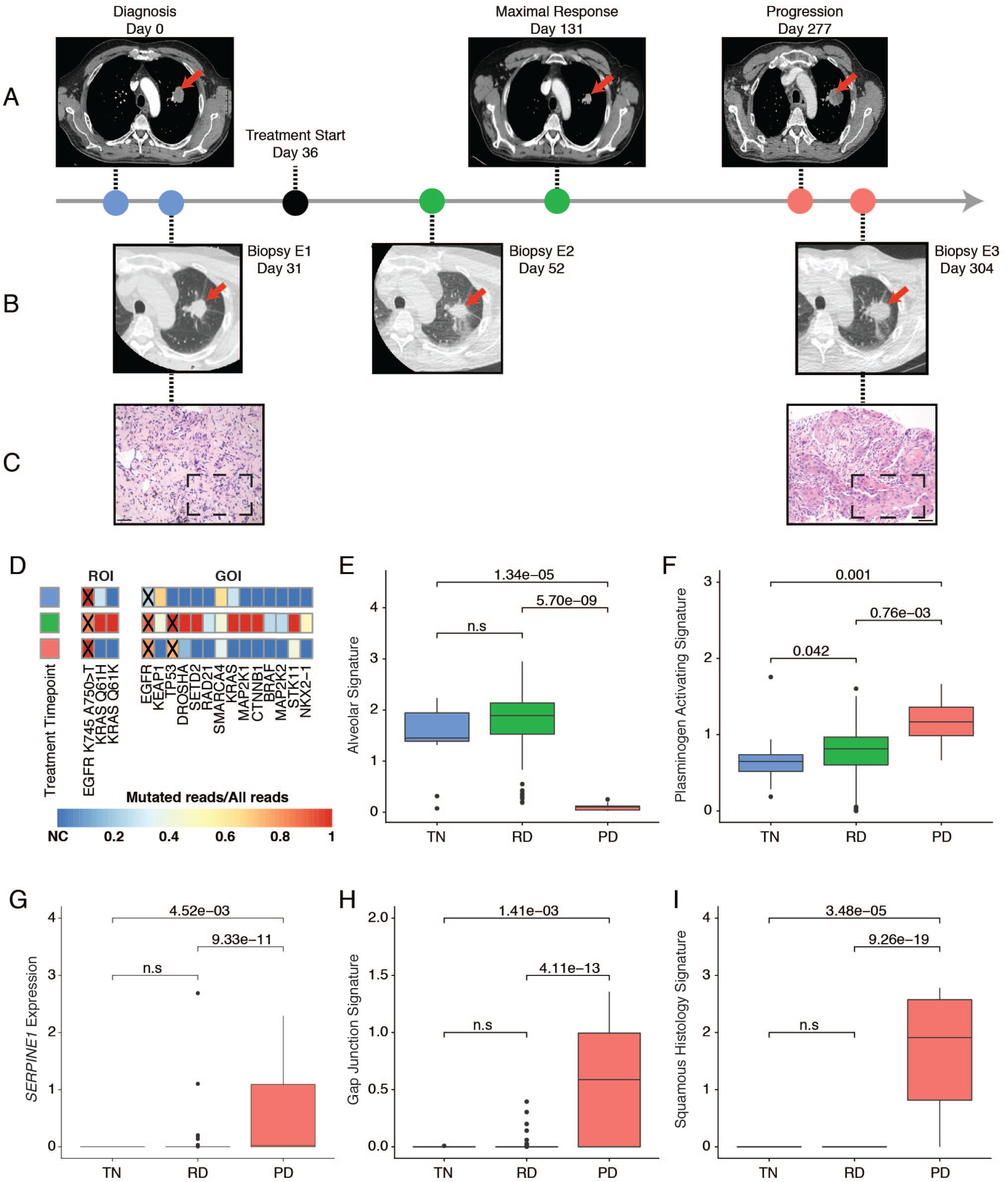
(A, B, C) Longitudinal timeline of patient treatment (A) Chest CT scan at each clinical evaluation timepoint (B) Biopsy timepoint with procedural CT scan (C) HE from treatment naïve and progression timepoints demonstrating adenocarcinoma and squamous cell carcinoma, respectively, scale bar indicates 50 µm (D) Heatmap of mutation state in COSMIC tier 1 mutated genes, where color represents mutated reads/all reads across treatment. Mutated genes as identified by clinical NGS are indicated by “X” (E-I) Boxplots of pathway signature changes (alveolar, plasminogen activating, SERPINE1, gap junction and squamous histology, respectively) across treatment timepoints.

**Fig S5 related to Figure 4.**
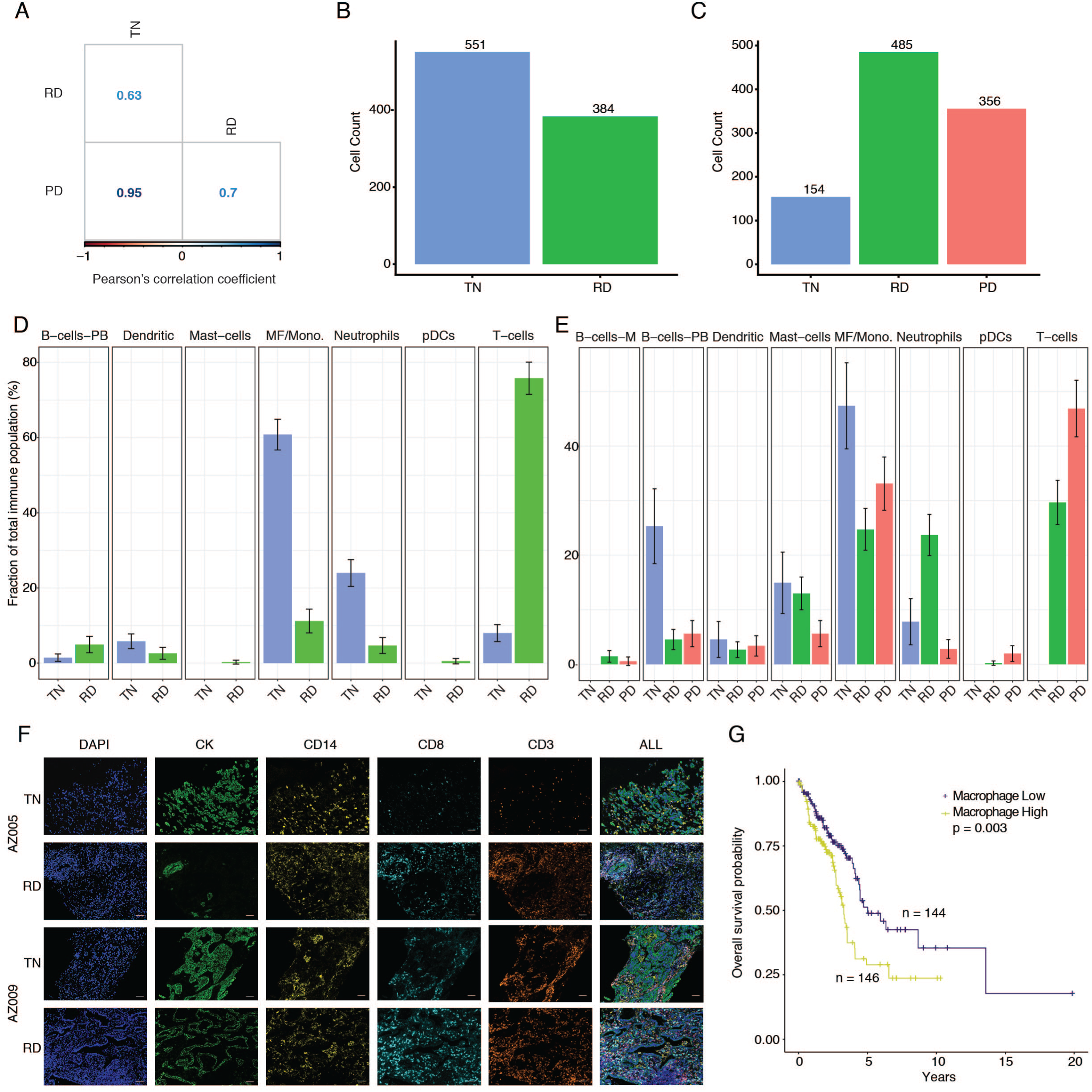
(A) Pairwise Pearson correlations between each treatment group’s immune cell compositions which corresponds to the fraction of each immune cell type’s abundance in the total immune cell population (B) Total immune cells for each biopsy of patient TH266. (C) Total immune cells for each biopsy of patient TH226. (D) Fraction of each immune cell sub-type for the two biopsies of patient TH266. Error bars indicate the 95% confidence interval for the calculated relative frequencies. (E) Fraction of each immune cell sub-type for the three biopsies of patient TH226. Error bars indicate the 95% confidence interval for the calculated relative frequencies. (F) Representative in situ immunofluorescence images from two patients with matched samples at different treatment timepoints, demonstrating fractional changes in the immune populations of macrophages and T-cells. Scale bars correspond to 50 µm (G) Kaplan-Meier plot of deconvoluted TCGA lung adenocarcinoma data showing the relation between OS and the fraction of macrophages for each patient. Patients were stratified by high and low macrophage fraction.

**Fig S6 related to Figure 5.**
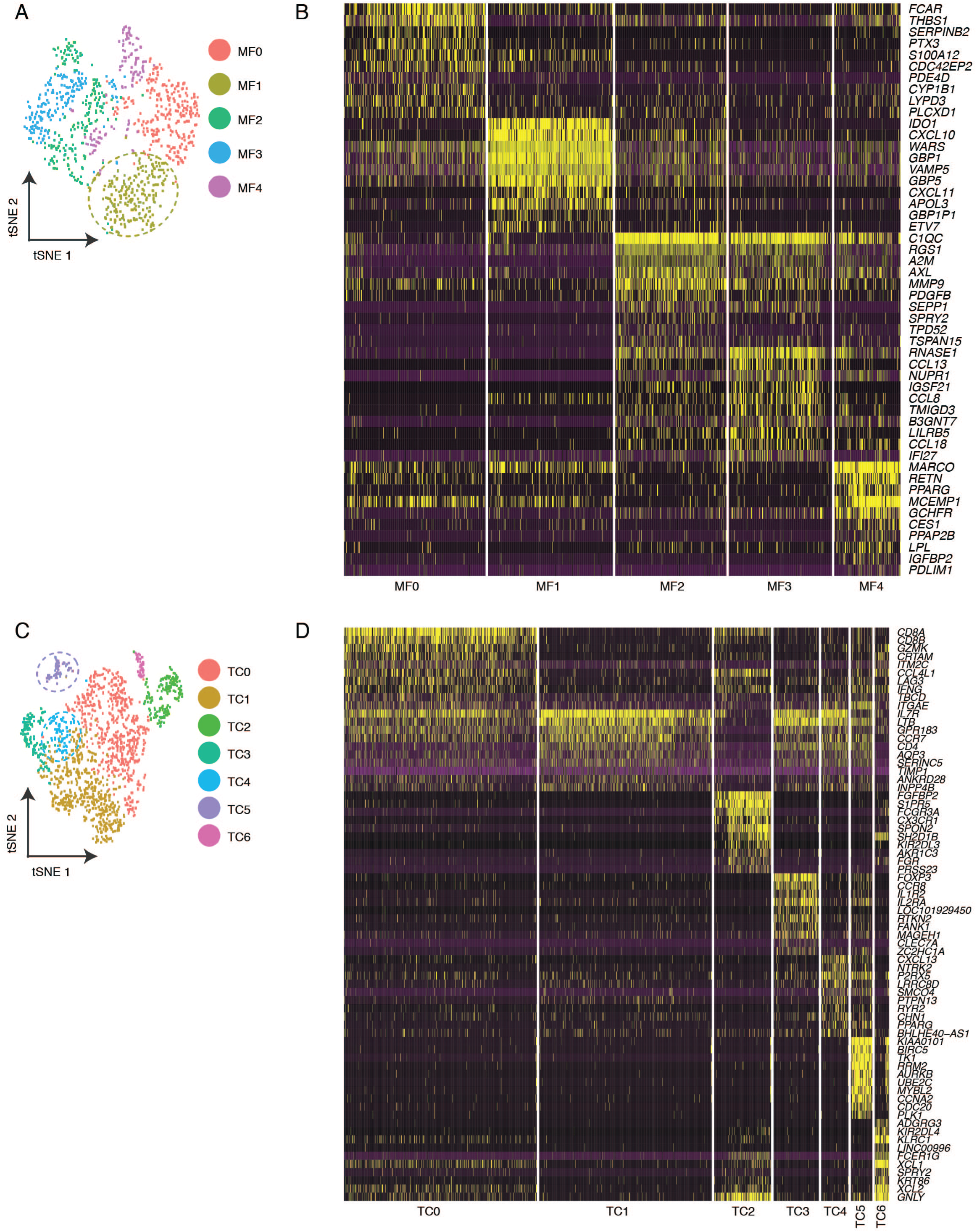
(A) t-SNE plot of all lung-derived macrophage cells. The cluster of cells enriched at PD (MF1) is highlighted (B) Heatmap showing the expression level of the top 10 differentially expressed genes for each macrophage clustert-SNE plot of all lung-derived T-cells. The clusters of cells enriched at PD (TC4 and TC5) are highlighted (D) Heatmap showing the expression level of the top 10 differentially expressed genes for each T-cell cluster.

